# A census of anti-CRISPR proteins reveals AcrIE9 as an inhibitor of *Escherichia coli* K12 Type IE CRISPR-Cas system

**DOI:** 10.1101/2025.05.07.652737

**Authors:** Dmitry Taranenko, Oksana Kotovskaya, Konstantin Kuznedelov, Daria Yanovskaya, Alina Demkina, Sofya Fardeeva, Viktor Mamontov, Kaiya Vierra, Nathaniel Burman, Dan Li, Minggui Wang, Blake Wiedenheft, Konstantin Severinov, Ekaterina Semenova, Artem Isaev

## Abstract

CRISPR-Cas adaptive immunity systems provide defense against mobile genetic elements and are often countered by diverse anti-CRISPR (Acr) proteins. The Type IE CRISPR-Cas of *Escherichia coli* K12 has been a model for structural and functional studies and is a part of the species’ core genome. However, this system is transcriptionally silent, which has fueled questions about its true biological function. To clarify the role of this system in defense, we carried out a census of Acr proteins found in *Enterobacterales* and identified AcrIE9 as a potent inhibitor of the *E. coli* K12 Type IE CRISPR-Cas system. While sharing little sequence identity, AcrIE9 proteins from *Pseudomonas* and *Escherichia* both interact with the Cas7 subunit of the Cascade complex, thus preventing its binding to DNA. We further show that AcrIE9 is genetically linked to AcrIE10, forming the most widespread anti-CRISPR cluster in *Enterobacterales*, and this module often co-occurs with a novel HTH-like protein with unusual architecture.

## Introduction

CRISPR-Cas (Clustered Regularly Interspaced Short Palindromic Repeats and CRISPR-associated genes) is one of the most widespread prokaryotic immunity systems, playing an important role in the co-evolution of phage and bacteria^1–3^. CRISPR-Cas systems utilize an adaptive immunity principle: at the adaptation stage, small DNA fragments derived from mobile genetic elements (MGE) are introduced into the CRISPR array via the action of conserved Cas1-Cas2 adaptation module^4,5^. Transcription of the CRISPR array and processing of the resulting non-coding RNA generates short guide CRISPR RNA molecules (crRNA)^6,7^, which interact with the effector Cas proteins, and mediate degradation of nucleic acid targets complementary to crRNA spacer sequences during the CRISPR interference stage^8,9^. CRISPR-Cas systems are divided into Class 1 with multi-protein effector complexes (e.g., Cascade and Cas3 in Type I systems), and Class 2 with single-protein effectors (e.g., Cas9, Cas12, Cas13)^10^. Based on the composition of the interference module, CRISPR-Cas systems are further classified into seven types and multiple subtypes^10,11^.

MGEs can evade CRISPR interference through accumulation of mutations in the targeted DNA regions, called protospacers^12^, mutations in the protospacer adjacent motif (PAM)^13^, or through activity of anti-CRISPR proteins (Acrs)^14–17^ and RNAs (Racrs)^18,19^. The mechanisms of Acr-mediated inhibition are quite diverse: from prevention of crRNA binding^18,20^, DNA mimicry^21–23^, inhibition of DNA recognition or cleavage by effectors^24–26^, to Cas protein modifications^27^, inactivation through dimerization^28^ or effector complexes disassembly^29^. As a countermeasure, some CRISPR-Cas systems with transcriptional auto-repression circuits could increase Cas protein expression in response to inhibition^30,31^. Acrs have been identified for the majority of CRISPR-Cas types and are found in diverse MGEs (lytic and temperate phages, conjugative plasmids, ICEs)^32–35^, where they form so-called anti-defense clusters regulated by Anti-CRISPR-associated (Aca) proteins that function as transcriptional repressors^32,36^. The presence of Acrs in mobile elements could be linked with the spread of antibiotic-resistance genes in clinical isolates of *Klebsiella*^37,38^ and other ESKAPE pathogens^39^.

Most *Escherichia coli* encode Type IE CRISPR-Cas system (*Ec*-IE) that can be associated with one or two cognate CRISPR arrays, while a few strains contain the Type IF (*Ec*-IF) that can be associated with two other CRISPR arrays^40,41^. The *Ec*-IE system served as a model for determining the principles of adaptation, crRNA maturation, and interference at the dawn of CRISPR-Cas research^42–46^. However, the activity of the *Ec*-IE system was either studied *in vitro* or in *E. coli* overexpression systems, because it is naturally transcriptionally silenced by H-NS^47,48^. This is a common bacterial strategy for the suppression of horizontally acquired xenogenic mobile elemetns^49,50^. Other transcriptional factors, such as LeuO, CRP, StpA, LrhA, and SspA contribute to the regulation of *cas* promoters^48,51–55^. Transcriptional silencing and the lack of spacers matching genomes of lytic phages, have led to a hypothesis that the *Ec*-IE system does not play a major role in anti-viral immunity of the host^56,57^. On the other hand, *Ec*-IE is present in ∼70% of sequenced *E. coli* isolates and thus represents a core part of the species pangenome^58,59^. It was speculated that *Ec*-IE can be activated under specific conditions, when transcriptional silencing is relieved^53^. The recent census of *Ec*-IE spacers in the gut microbiome revealed preferential targeting of prophages^60^, while independent work proposed the role of *Ec*- IE in control of cryptic prophages^61^.

Existence of Acrs active against *Ec*-IE can support a role for this system in genetic conflicts with MGEs in natural populations. Machine learning-based methods have predicted multiple Acr candidates within mobile elements of *E. coli*^62–64^. This could be attributed to the overrepresentation of *E. coli* genomes in the databases or may suggest a hidden diversity of Acrs active against the *Ec*-IE system. Earlier experimental studies have not identified Acrs against the *Ec*-IE system^65^, while more recent works demonstrated anti-*Ec*-IE activity of the AcrIE10^38^ and AcrIC6*^30^ proteins. Nevertheless, a dedicated study of the *Ec*-IE sensitivity to Acrs has not been carried out.

Here, we analyzed the distribution of Acrs within mobile elements of *E. coli* and *Enterobacterales* and experimentally investigated a collection of such Acrs, along with selected Acrs derived from *Pseudomonas aeruginosa*. We demonstrate that *Ec*_AcrIE9 and *Pa*_AcrIE9 robustly inhibit the *Ec*-IE system, while *Ec*_AcrIF15 and *Kp*_AcrIF22 exhibit moderate inhibitory activity against the *Ec*-IF system. We further show that AcrIE9 binds to the Cas7 subunit of Cascade, inhibiting its interaction with target DNA. Analysis of the genomic loci encoding AcrIE9 revealed a strong functional link with the AcrIE10 protein and a novel HTH-like protein with unusual architecture. These results provide the first mechanistic insight into Acr activity against the model Type IE CRISPR-Cas of *E. coli* K12 and have implications for the co-evolution of CRISPR-Cas and MGE in pathogenic bacteria.

## Results

### Multiple Acr classes are found in *Enterobacterales* MGEs

To identify Acr proteins encoded by *Enterobacterales* and their MGEs, we leveraged the dbAPIS database of anti-immune proteins^35^ and performed an Acr HMM profile search against prokaryotic and viral protein sequences derived from non-redundant database (nr BLAST) of microbial and MGE genomes (**Extended Data File 1**). **Figure 1A** sums up the phylogenetic distribution of Acr classes, for which at least one hit was found within *Enterobacterales* genomes or within phages infecting *Enterobacterales* hosts. These data reveal that AcrIE8, IE9, IF11, IF15-22, IIA7, and VA2 are predominantly found within *Enterobacterales*. Genus-level analysis further demonstrates a “specialization” of different Acr classes towards specific bacteria, and multiple AcrIE and AcrIF homologs were found within *Escherichia* genomes (**Figure 1B**). In this dataset, AcrIE9 represents the most abundant Acr variant found in more than 54,000 instances. It should be noted that AcrIE10^38^ was not present in the dbAPIS HMM profiles database, while our downstream analysis identifies its comparable abundance to AcrIE9. Next, we focused on the genomic environment of Acrs within *Enterobacterales* and found that the majority of Acr genes are encoded by MGEs: phages (including metagenomic viral contigs), prophages, plasmids, and conjugative plasmids (**Figure 1C**). We selected 9 candidates for experimental validation: 2 proteins predicted to be active against IE system, and 7 anti-IF proteins (**Figure 1A, Supplementary Table 1**). To balance the collection, we selected homologs of Acr proteins from *P*.*aeruginosa* as additional controls (2 anti-IE and 4 anti-IF). Since multiple Acrs were shown to possess broad spectrum activity against different types of CRISPR-Cas systems, including crossinhibitory activity against IE and IF subtypes^23^, all candidates were tested against the *Ec*-IE and the *Ec*-IF systems.

**Figure 1:**
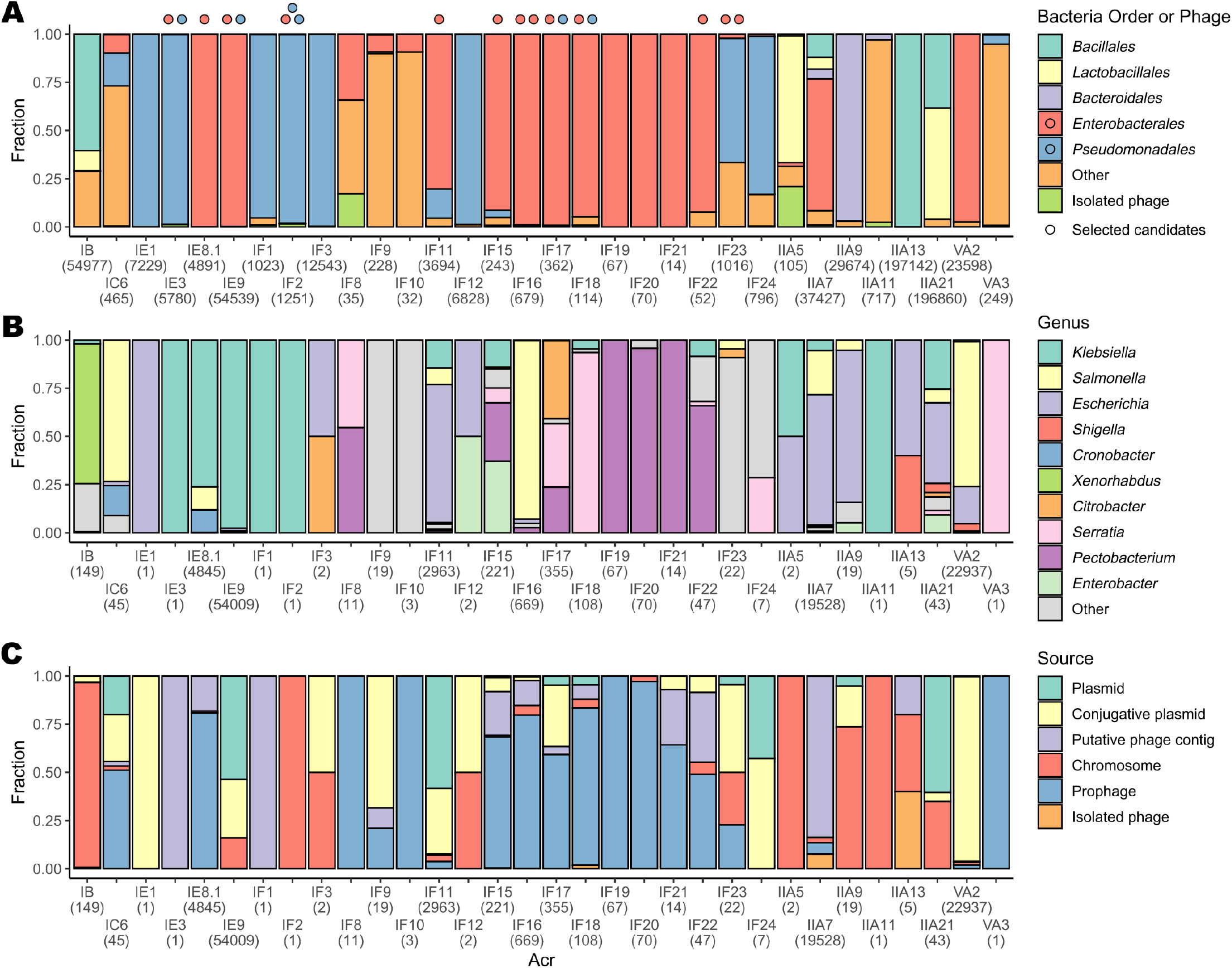
Acr classes distribution among *Enterobacterales*. **(A)** Distribution of known anti-CRISPR proteins among phage genomes and different bacterial groups. Colored circles at the top represent selected candidates for experimental evaluation, red – from *Enterobacterales*, blue – from *Pseudomonadales*. **(B)** Genus level taxonomic distribution of Acrs found in *Enterobacterales*. **(C)** Local genomic context of Acr genes found in *Enterobacterales*, as predicted by geNomad. `Putative phage contig` category includes predicted phage contigs lacking terminal repeats. `Chromosome` category includes all loci, not found to be associated with other categories. Numbers in parenthesis for all panels represent number of loci found.

### AcrIE9 from *E. coli* and *P. aeruginosa* inhibits Type IE CRISPR-Cas of *E. coli*

To estimate the anti-CRISPR activity against the *Ec*-IE system, we used F’ KD263 strain^66^, encoding a G8 spacer targeting M13 phage and chromosomal *cas* genes under the control of the arabinose- and IPTG-inducible promoters (**Figure 2A**). Selected Acr candidates were synthesized, cloned, and expressed from a pBAD plasmid under the control of an *araBAD* promoter. While in the presence of an empty pBAD vector (EV), the induced *Ec*-IE system provided robust defence against M13 in phage plaque assay, expression of the *Pa*_AcrIE9 and *Ec*_AcrIE9 restored phage plaquing efficiency (**Figure 2B**). To further validate this effect, we used the SS80 strain, a derivative of the KD263 with a spacer against lytic phage T7^44^ (**Figure 2C**). Again, *Pa*_AcrIE9 and *Ec*_AcrIE9 completely suppress the induced *Ec*-IE immune system in both phage plating assay and liquid culture infection (**Figure 2D, Supplementary Figure 1A-B**). However, at low Multiplicity of Infection (MOI) *Pa*_AcrIE9 outperforms *Ec*_AcrIE9 (**Supplementary Figure 1C**). Lastly, we investigated Acr activity in conditions of CRISPR-Cas expression from native promoters in *Δhns* background, since overexpression of Cas proteins from inducible promoters could mask inhibitory effects of some Acrs. We used BW39671 *Δhns* host^45^ encoding λT3 spacer targeting phage λ (**Figure 2F**) and estimated phage *λ*_*vir*_ plaquing efficiency. Cells with a non-targeting spacer were used as a negative control, and showed a lack of defence. While the inhibitory effect of *Pa*_AcrIE9 and *Ec*_AcrIE9 on the *Ec*-IE system was confirmed, no other Acr protein demonstrated robust inhibition (**Figure 2F**). We additionally monitored the accumulation of the phage particles in the spent liquid media of BW39671 *Δhns* cells infected with *λ*_*vir*_. In line with the EOP assay, *Pa*_AcrIE9 or *Ec*_AcrIE9 expression enhanced accumulation of phage progeny, although did not completely inhibit the native CRISPR-Cas system (**Supplementary Figure 1D,E**).

**Figure 2:**
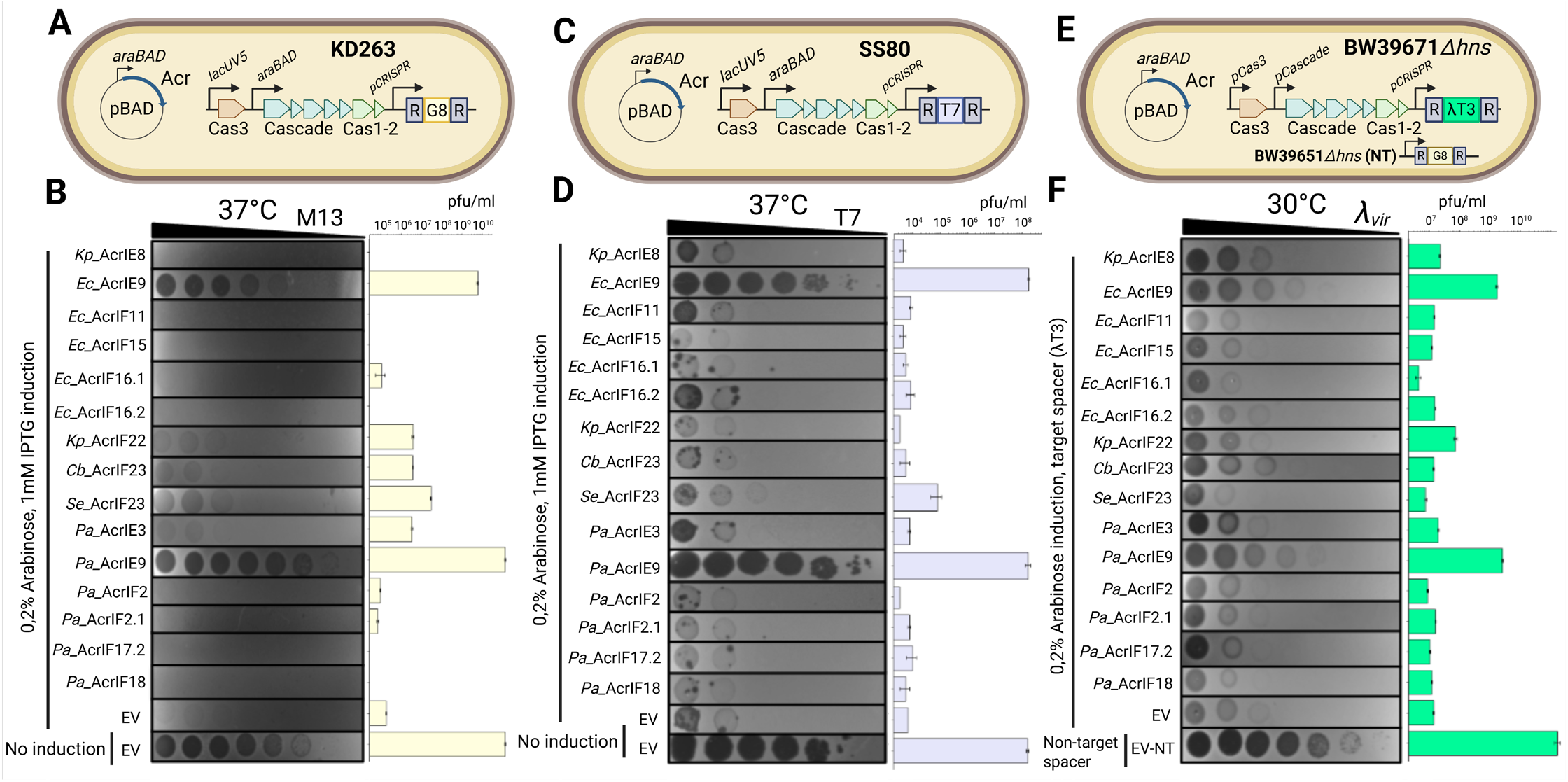
Screening of Acr candidates reveals anti-CRISPR proteins active against Type I-E CRISPR-Cas of *E*.*coli*. **(A)** *E. coli* KD263 strain with a M13 phage-targeting spacer G8 was used for M13 phage infection assay. **(B)** EOP with phage M13 on the lawns of KD263 cells with induced expression of Cas proteins and Acr candidates. Cultures with empty pBAD vector and without induction of CRISPR-Cas expression were used as controls. **(C)** *E. coli* SS80 strain, a KD263 derivative with T7-targeting spacer, was used for T7 infection assay. **(D)** EOP with phage T7 on the lawns of SS80 cells with induced expression of Cas proteins and Acr candidates. Cultures with empty pBAD vector and without induction of CRISPR-Cas expression were used as controls. **(E)** *E. coli* BW39671 strain, a BW25113 *Δhns* derivative with a *λ* phage-targeting spacer T3, was used for *λ*_*vir*_ phage infection assay. **(F)** EOP with phage *λ*_*vir*_ on the lawns of BW39671 cells with induced expression of Acr candidates. Cultures without a *λ*_*vir*_ targeting spacer and with empty pBAD vector were used as controls. EOPs presented on panel **(B)** and **(D)** were performed at 37C, while EOP from the panel **(F)** was performed at 30C. Acr expression was induced with 0.2% L-arabinose, CRISPR-Cas expression at the EOPs presented on panel **(B)** and **(D)** was induced with 0.2% L-arabinose and 1 mM IPTG, while EOP from the panel **(F)** was performed at conditions of native CRISPR-Cas expression. All experiments were performed in biological triplicates and bars on the right represent mean pfu/ml values.

A recently published ML-based model predicted multiple novel Acr classes and *E. coli* represented the largest source of identified canidates^62^. To test the activity of these candidates we selected, synthesized, and cloned representatives from the top 13 anti-IEs found in *E. coli* (**Supplementary Table 2**). While the two candidates representing Acr candidate clusters 2320 and 1797, exhibited high toxicity to the host, the expression of the remaining candidates neither resulted in toxic phenotypes nor showed notable *Ec*-IE suppression (**Supplementary Figure 2A,B**).

### Acr proteins provide moderate anti-CRISPR activity against Type IF CRISPR-Cas of E. coli LF82

Although not as widespread as Type IE, Type IF CRISPR-Cas system can be found in many *E. coli* isolates^40,41^. Previous work reported that the chromosomally encoded *Ec*-IF system from LF82 strain interferes with the plasmid stability^67^. Despite multiple attempts, we were unable to demonstrate LF82 *Ec*-IF system interference against M13 phage infection or plasmid transformation under native conditions. Thus, we turned to the plasmid-mediated expression of the *Ec*-IF LF82 system in a CRISPR-deficient BB101 *E. coli* host^68^ (**Figure 3A**). In the presence of the targeting spacer, the *Ec*-IF LF82 system reduced the phage Mu plaquing ability by two orders of magnitude (**Figure 3B**). Expression of *Ec*_AcrIF15, *Kp*_AcrIF22, and *Pa*_AcrIF18 provides moderate (i.e., 10x) inhibition of the *Ec*-IF LF82 interference, supporting the existence of active anti-IF Acrs in MGEs of *Entrobacterales* (**Figure 3B**). ML-predicted clusters lacked an activity against the *Ec*-IF system (**Supplementary Figure 2C,D**). Given the relatively low Acr activity against the IF system, we focused on elucidating the mechanisms of *Ec*-IE inhibition by AcrIE9 in subsequent analyses.

**Figure 3:**
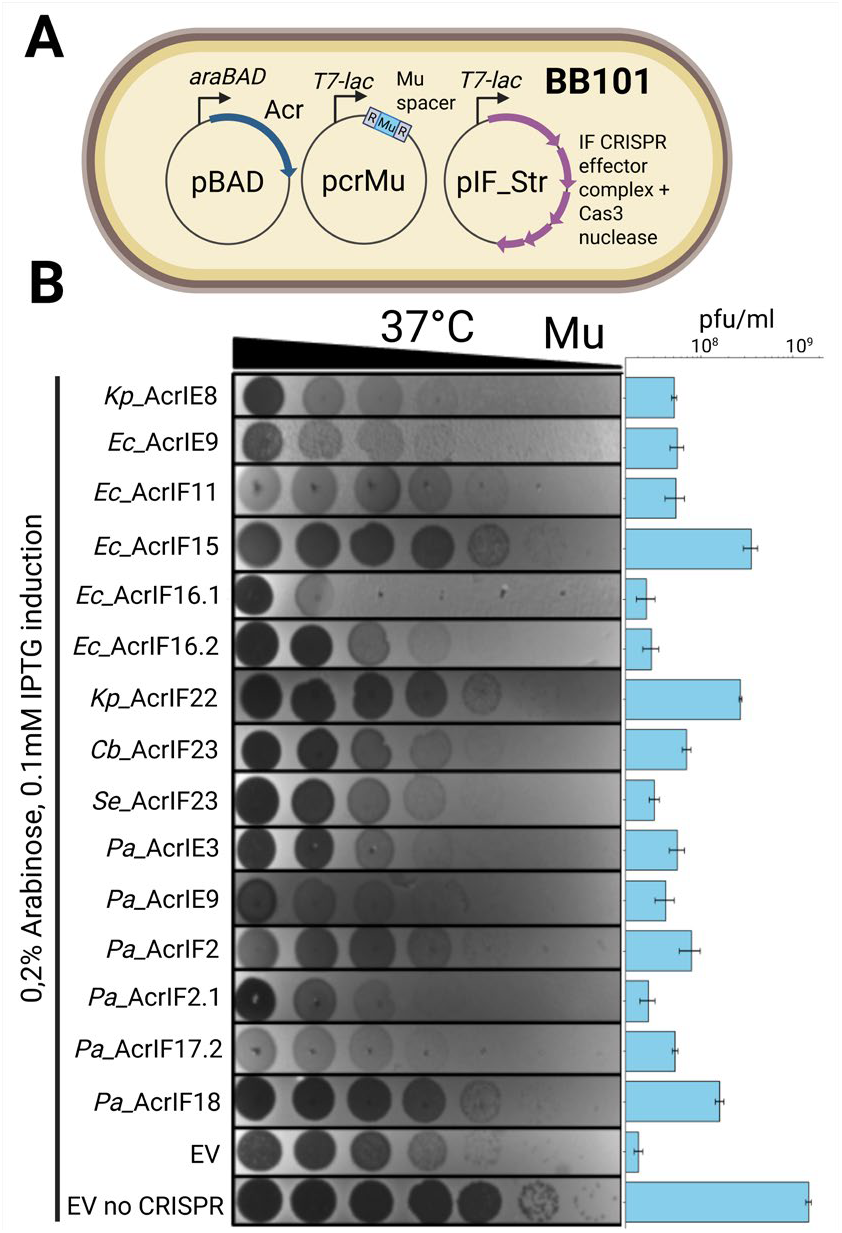
Screening of Acr candidates reveals potential anti-CRISPR proteins targeting I-F CRISPR of *E*.*coli* LF82. **(A)** *E. coli* BB101 strain with a plasmid-encoded IF LF82 system and phage-targeting spacer was used for the phage Mu phage infection assay. **(B)** EOP with phage Mu on the lawns of BB101 cells with induced expression of Cas proteins and Acr candidates. Cultures with empty pBAD vector and without CRISPR-Cas encoding plasmid were used as controls. Acr expression was induced with 0.2% L-arabinose, CRISPR-Cas and spacer expression was induced with 0.1 mM IPTG. All experiments were performed in biological triplicates and bars on the right represent mean pfu/ml values.

### AcrIE9 inhibits primed, but not naïve, adaptation

We started with testing whether AcrIE9 inhibits the incorporation of new spacers into the CRISPR array. For this, we overexpressed Cas1-Cas2 proteins in the *E. coli* BL21-AI host carrying CRISPR array but lacking other Cas proteins. We then monitored CRISPR array expansion via PCR in the presence and absence of AcrIE9 (**Figure 4A**). After 24h of cultivation, the CRISPR array expanded for one or two new spacers, and spacer acquisition remained unaffected by the presence of AcrIE9 (**Figure 4B**).

**Figure 4:**
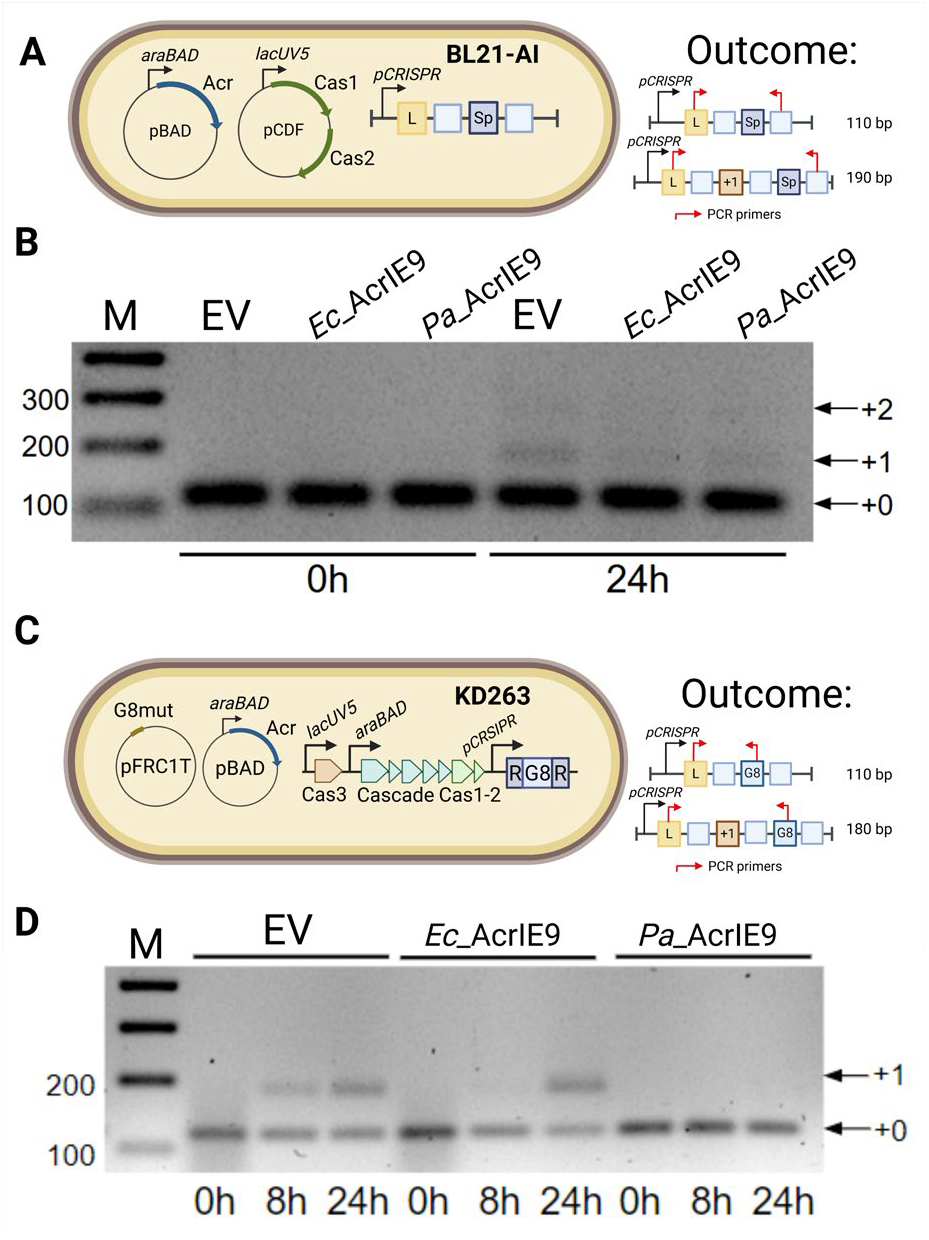
AcrIE9 inhibits primed but not naïve adaptation. **(A)** *E. coli* BL21-AI strain with a plasmid-encoded Cas1-Cas2 was used for the naïve adaptation assay. Acr expression was induced with 0.2% L-arabinose, Cas1-Cas2 expression was induced with 1 mM IPTG. **(B)** Agarose gel representing CRISPR arrays after naïve adaptation. Expanded arrays are labeled with arrows. **(C)** *E. coli* KD263 strain transformed with a priming plasmid was used for the primed adaptation assay. Acr expression was induced with 0.2% L-arabinose, CRISPR-Cas expression was induced with 0.2% L-arabinose and 1 mM IPTG. **(D)** Agarose gel representing CRISPR arrays after primed adaptation. Expanded arrays are labeled with arrows. Representative gels of the experiments carrying in biological triplicates are shown on the panels **(B)** and **(D)**. Expected size of the products after PCR with CRISPR array-specific primers with non-expanded and expanded array templates are shown on the right on the panels **(A)** and **(C)**.

Some CRISPR-Cas systems, including *Ec*-IE, have evolved a primed adaptation mechanism, which facilitates efficient spacer acquisition from CRISPR-targeted DNA^69^. This process is triggered by a pre-existing spacer that matches a sequence within the genome of MGE. Primed adaptation efficiently restricts the accumulation of escapers. This mode of spacer acquisition relies on the coordinated action of both the interference and adaptation modules and is, therefore, strictly dependent on the presence of Cas interference proteins^69^. To investigate the effect of AcrIE9 on primed adaptation, we transformed *E. coli* KD263 with a targeted plasmid, carrying a single mismatch (C1T) in the protospacer region complementary to the G8 spacer^70^ (**Figure 4C**). In the absence of AcrIE9, expansion of the CRISPR array was already observed 8h after *Ec*-IE induction (**Figure 4D**). In contrast, expression of *Ec*_AcrIE9 resulted in CRISPR array expansion only 24h after *Ec*-IE induction, whereas expression of *Pa*_AcrIE9 completely inhibited primed adaptation. This aligns with the more pronounced anti-CRISPR effect observed in liquid culture infection (**Supplementary Figure 1C**). Together, these results indicate that AcrIE9 does not affect naïve adaptation but specifically inhibits primed adaptation, which depends on the activity of the Cascade and Cas3.

### AcrIE9 binds to the Cas7 subunit and prevents Cascade interaction with DNA *in vivo* and *in vitro*

To identify the mechanism of CRISPR inhibition, we tested the ability of *Ec*-IE Cascade to bind DNA in the presence of AcrIE9. In Type I CRISPR-Cas systems, the Cascade complex is guided by crRNA to a complementary DNA target. Binding of the target DNA leads to the formation of an R-loop, which induces conformational changes required for the recruitment of the processive nuclease-helicase Cas3^71,72^. In the absence of Cas3, Cascade remains bound to the protospacer and can act as a transcriptional silencer^73,74^. This property allows for the discrimination of Acr mechanisms (i.e., blocking DNA degradation versus inhibition of Cascade assembly, or target DNA recognition). We used *Δcas3* KD454 strain with inducible *Ec*-IE system^75^ and introduced a G8 protospacer in the promoter of sfGFP gene expressed from a plasmid (**Figure 5A**). As expected, expression of sfGFP was repressed in conditions of CRISPR-Cas induction and the presence of the targeted G8 protospacer (**Figure 5B**). Expression of *Ec*_AcrIE9 and *Pa*_AcrIE9 slightly reduced cell growth, indicating intermediate toxicity, yet restored sfGFP production. This suggests that AcrIE9 prevents DNA recognition by the *Ec*-IE Cascade Complex (**Figure 5B, Supplementary Figure 3**).

**Figure 5:**
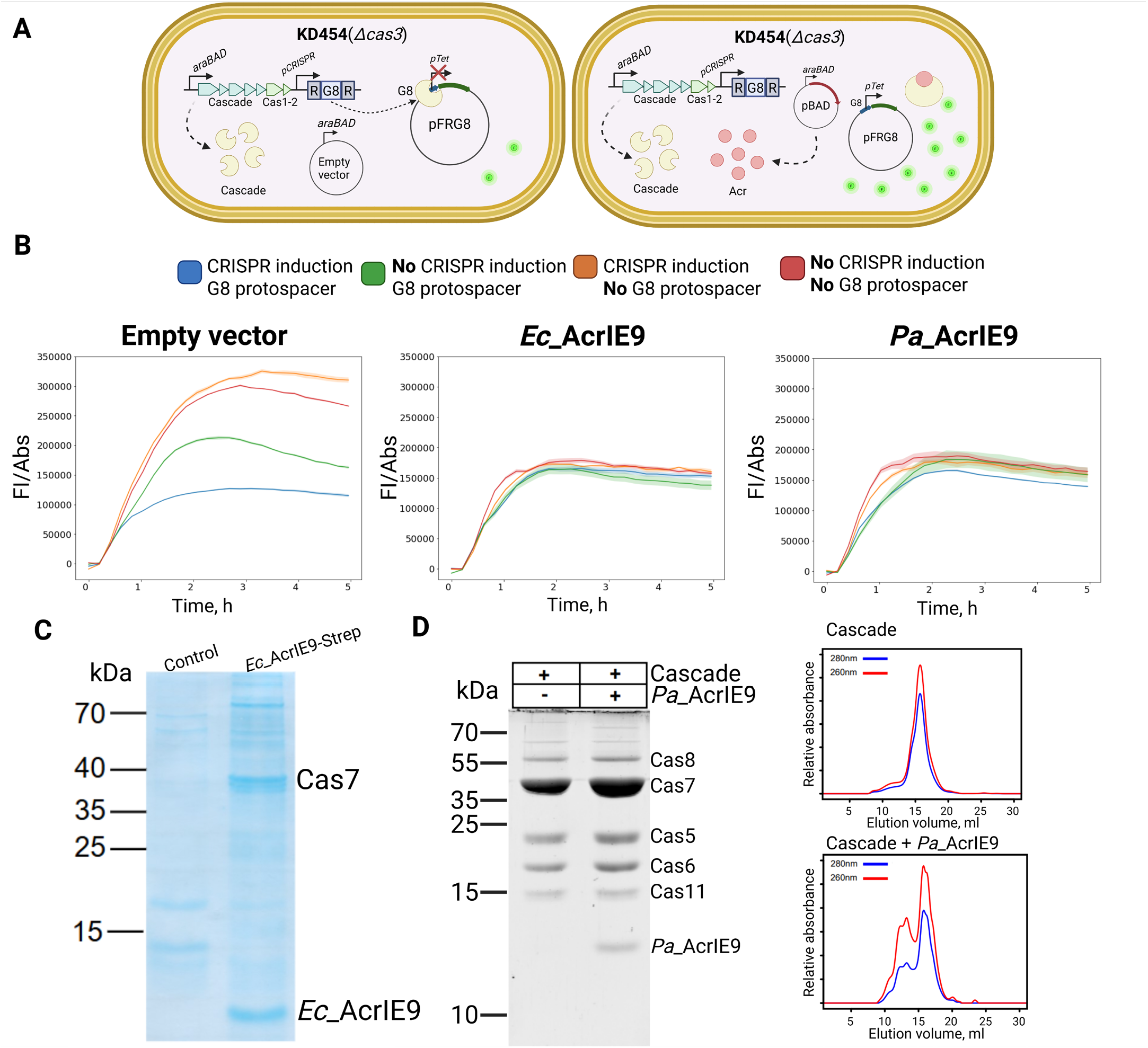
AcrIE9 targets Cas7 subunit of Cascade *in vivo* and interferes with Cascade DNA binding. **(A)** *E. coli* KD454 strain, a KD263 derivative lacking Cas3, was used for CRISPR transcriptional silencing assay. sfGFP was expressed from *pTet* containing or lacking G8 protospacer. Acr expression was induced with 0.2% L-arabinose, CRISPR-Cas expression was induced with 0.2% L-arabinose and 1 mM IPTG, sfGFP expression was induced with 200 ng/mL aTc. A scheme of the assay and conditions color code is shown on the right. **(B)** Dynamics of sfGFP production in KD454 cells carrying an empty pBAD vector or expressing *Ec*_AcrIE9/*Pa*_AcrIE9. Fluorescence intensity was normalized to the culture optical density. **(C)** *In vivo* pull-down of *Ec*_AcrIE9 with a C-terminal Strep tag in *E*.*coli* KD263 after induction of untagged *Ec*_IE Cascade expression. Control represents a similar experiment carried with AcrIE9_Ec without the Strep tag. Identity of the labeled proteins on was determined via MALDI-TOF mass-spectrometry. **(D)** SDS-PAGE analysis of Cascade purified from cell cultures with or without *Pa*_AcrIE9 co-expression (left panel). SEC elution profiles of Cascade and Cascade:*Pa*_AcrIE9 complexes run on Superose 6 10/300 column (right panel).

To check if AcrIE9 directly interacts with Cascade, we constructed a C-terminally Strep-tagged version of *Ec*_AcrIE9 and performed an *in vivo* pull-down experiment in KD263 cells producing all *Ec*_IE subunits. Cells expressing untagged AcrIE9 were used as a negative control. A coomassie stained SDSPAGE of the pulldown reveals Cas7 subunit as the major AcrIE9 binding partner (**Figure 5C**). To clarify whether AcrIE9 binds to the free Cas7 or the assembled Cascade complex, we performed a total proteomic analysis of the StrepTrap column eluates that additionally revealed Cas11 and Cas8e among the most enriched proteins (**Supplementary Figure 4A**). The lack of other Cascade subunits could reflect they lower abundance and/or decreased stability of the protein complexes not directly bound to a bait (AcrIE9). A similar analysis performed with *Pa*_AcrIE9 also confirmed co-purification of the Cas7 and Cas8 subunits, despite a weaker binding of the bait protein to the resin (**Supplementary Figure 4B**). We further performed purification of Strep-tag labeled *Ec*_IE Cascade from cells coexpressing untagged *Pa*_AcrIE9 and via SEC and SDS-PAGE analysis demonstrated the formation of a stable complex, containing all expected Cascade subunits (**Figure 5D**). Together, the results confirm direct interaction of AcrIE9 with complete Cascade complex *in vivo*.

To validate these results, we purified AcrIE9 and studied its interaction with Cascade *in vitro*. Since *Pa*_AcrIE9 was more stable after purification, most downstream experiments were performed with this variant. First, we noticed that AcrIE9 on it’s own has double-stranded but not single-stranded DNA binding activity (**Figure 6A**). Incubation with DNA binding competitor heparin reduces gel shift (**Supplementary Figure 5A)**. In addition, protein staining after native PAGE revealed that irrelevant ot the presence of DNA AcrIE9 migrates as a higher molecular weight complex, while incubation with heparin reduces its size (**Figure 6A, Supplementary Figure 5A**). Since Acr proteins can inhibit CRISPR-Cas DNA cleavage activity through effects on DNA topology^76^, we studied whether AcrIE9 can provide a non-sequence specific protection against nucleases. Excluding this hypothesis, plasmid DNA remained sensitive to Cas9 cleavage even in the presence of high concentrations of AcrIE9 (**Supplementary Figure 5B**).

**Figure 6:**
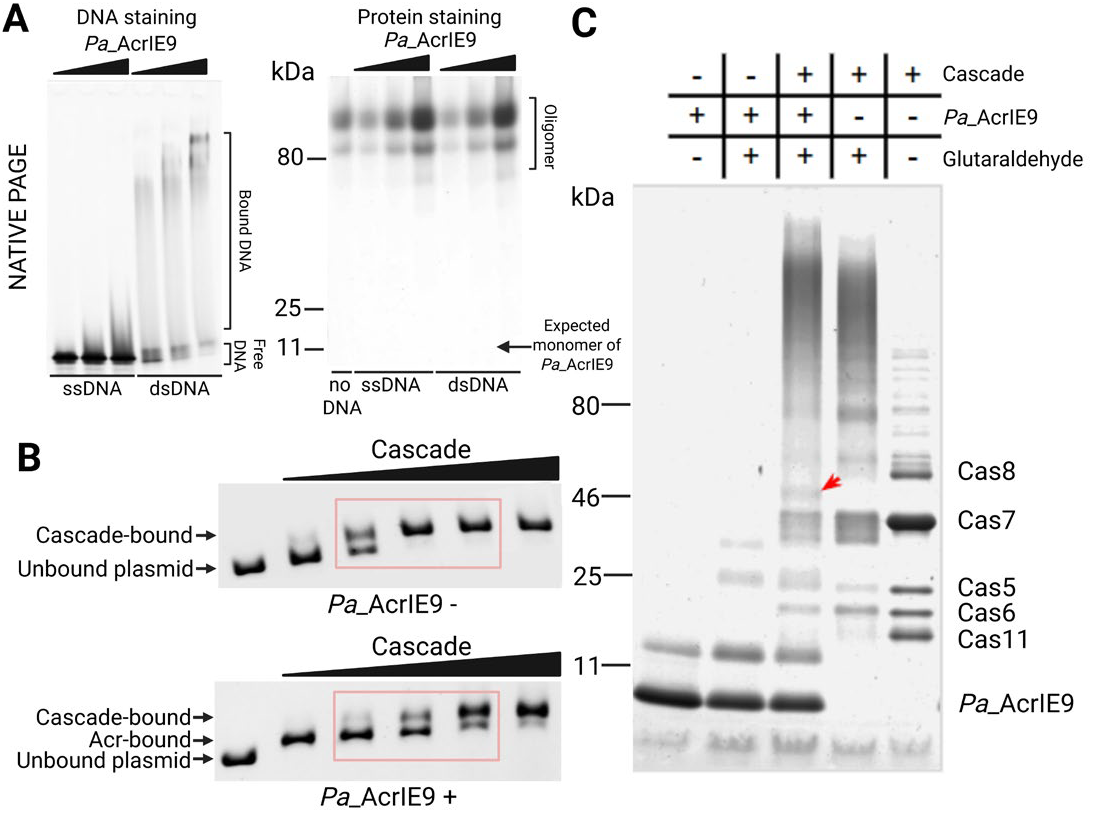
AcrIE9 binds to the Cas7 subunit of the assembled Cascade complex and interferes with target DNA recognition *in vitro*. **(A)** EMSA demonstrates *Pa*_AcrIE9 binding to dsDNA. 0.25 μM of 52-bp single-stranded or double-stranded DNA was incubated with *Pa*_AcrIE9 at increasing concentrations (17, 33 and 66 μM). **(B)** EMSA with increasing concentrations of *Ec*_IE Cascade (20 – 320 nM) and 4.5 nM of target plasmid in the presence or absence of an excess (1.5 μM) of *Pa*_AcrIE9. With-out AcrIE9, a complete plasmid shift is achieved (top panel). In the presence of *Pa*_AcrIE9, two gel shifts associated with consequent binding of the *Pa*_AcrIE9 and *Ec*_IE Cascade could be observed, and Cascade binding is incomplete (bottom panel). **(C)** *In vitro* glutaraldehyde (0.5%) crosslinking assay between 23 μM *Pa*_AcrIE9 and 6 μM of *Ec*_IE Cascade, resulting in the appearance of novel band migrating above Cas7 (red arrow), that was confirmed to represent Cas7-AcrIE9 adduct.

We further studied whether AcrIE9 inhibits *in vitro* interaction of *Ec*_IE Cascade with the target plasmid DNA. First, we established the range of Cascade concentrations that produce gel shift in the absence of AcrIE9 (**Figure 6B**). Next, we performed EMSA in the presence of an excess of AcrIE9. Here, two gel shifts can be observed, first associated with the DNA binding by AcrIE9, and the second shift mediated by Cascade binding (**Figure 6B**). However, the second shift required higher concentrations of Cascade, compared to conditions without AcrIE9. To prove that AcrIE9 inhibits Cascade DNA binding, we performed EMSA with a fixed concentration of Cascade and increasing concentrations of AcrIE9. This assay confirmed disruption of a complex with the target plasmid DNA, and comparable results were achieved for both, *Pa*_AcrIE9 and *Ec*_AcrIE9 (**Supplementary Figure 5C**).

*In vivo* data demonstrated that Cas7 subunit is the major AcrIE9 binding partner (**Figure 5C**). To confirm this observation *in vitro*, we performed cross-linking of the Cascade in the presence or absence of AcrIE9; a control reaction contained AcrIE9 only. When both Cascade and AcrIE9 were present in the cross-linking reaction, we observed a novel band migrating above Cas7 (**Figure 6C**). Mass-spectrometry analysis confirmed the presence of AcrIE9- and Cas7-derived peptides in this band, supporting the formation of a covalent adduct between two proteins. To investigate a possible role of other Cas subunits, we repeated cross-linking experiments with Cascade subcomplexes (Cas8-11-7-5-6, Cas11-7-5 and Cas11-7-5-6) at increasing concentrations of AcrIE9. Intensity of the fusion protein increased with AcrIE9 concentration, and this band was observed for all conditions tested, suggesting that Cas8 and Cas6 subunits are unlikely to contribute to AcrIE9 binding (**Supplementary Figure 5D**).

Together, *in vivo* and *in vitro* data support the model of AcrIE9 binding to the Cas7 subunit of Cascade, leading to compromised target DNA recognition.

### AcrIE9 can be harnessed for the control of Cas3-mediated genome editing

While single-subunit Class 2 CRISPR-Cas effectors become the standard genome editing tools, effectors of the Class 1 complexes, including Type IE Cascade+Cas3, also have promising applications. Cas3 is a processive nuclease/helicase and thus can be harnessed for the introduction of long chromosomal deletions^77–79^. Acr proteins have already proved useful for the temporal control of CRISPR-Cas activity and reduction of the off-target effects^80–82^, and AcrIE9 can potentially be used to control the activity of Cascade+Cas3 complexes. To investigate this possibility, we used *E. coli* KD504 strain, encoding a self-targeting spacer (**Figure 7A**).

**Figure 7:**
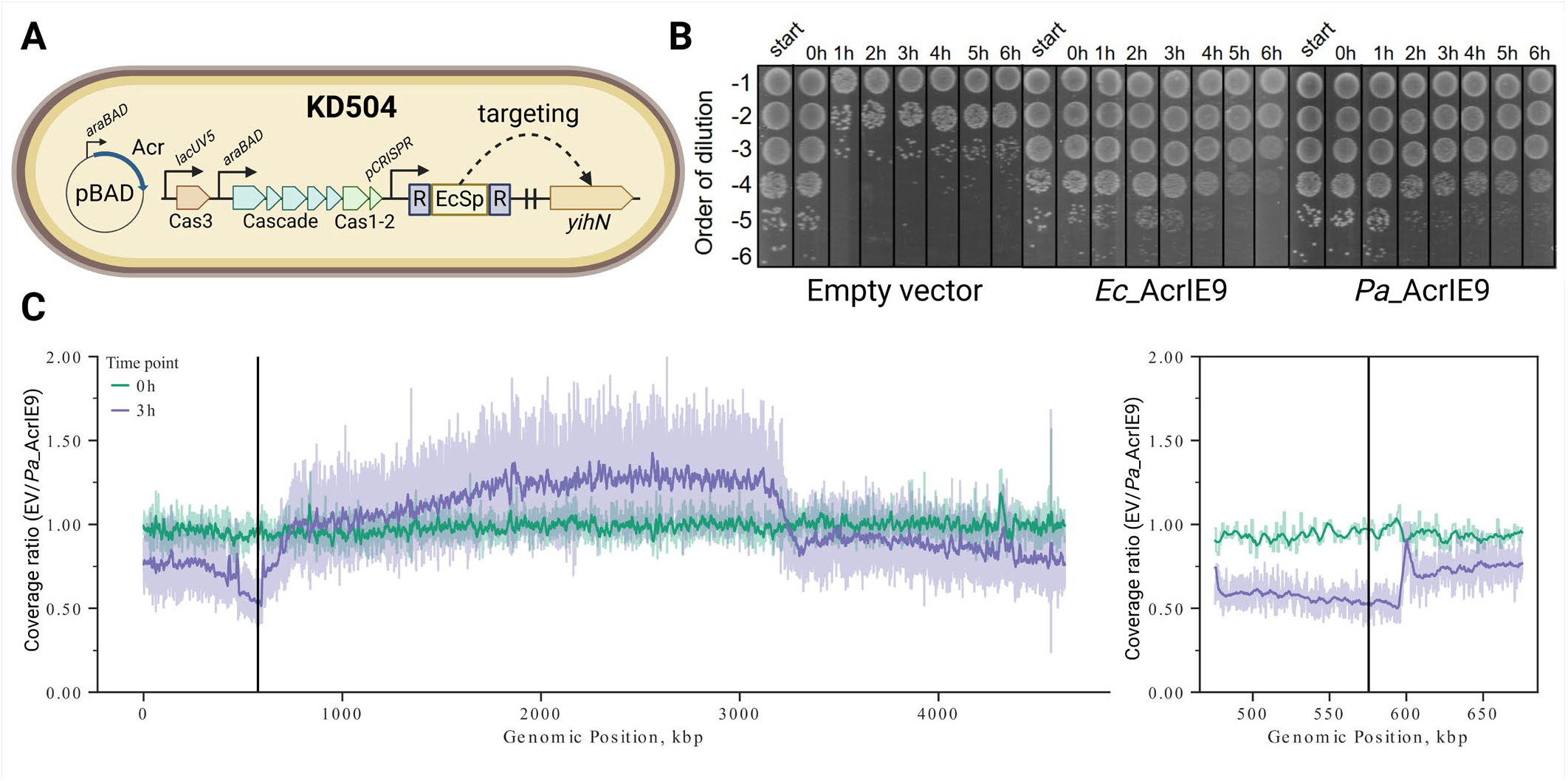
AcrIE9 prevents genomic DNA degradation in a self-targeting CRISPR-Cas system. **(A)** *E. coli* KD504 strain, a KD263 derivative encoding a self-targeting *yihN* spacer, was used for the CRISPR toxicity assay. Acr expression was induced with 0.2% L-arabinose, CRISPR-Cas expression was induced with 0.2% L-arabinose and 1 mM IPTG. **(B)** Cell toxicity after induction of CRISPR-Cas self-targeting. CRISPR-Cas was induced at time zero and at indicated time points cells were plated on M9 media supplemented with 0.2% glucose. Representative plates from an experiment performed in biological triplicates are presented. **(C)** Genomic DNA coverage in self-targeting KD504 cells after CRISPR-Cas induction in conditions of *Pa*_AcrIE9 expression or in the presence of the empty pBAD vector. Bold curves represent moving average for coverage ratios before and 3h after induction of CRISPR-Cas expression. Window size is 10kb for the left panel and 5kbp for the right panel zoomed in at the *yihN* protospacer region (black vertical line). Semi-transparent curves represent 1x-normalized coverage ratios for each nucleotide position.

Induction of *Ec*-IE system expression in this strain resulted in the rapid loss of cell viability, while the presence of *Ec*_AcrIE9 or *Pa*_AcrIE9 prevented cell death caused by the Cas3- mediated degradation of the host DNA (**Figure 7B**). We sequenced genomic DNA purified from cells with induced CRISPR self-targeting, and compared genome coverage obtained from cells containing empty pBAD plasmid (EV) to those expressing *Pa*_AcrIE9 (**Figure 7C**). The results reveal extensive degradation of the chromosomal DNA stretching up to 1.8 Mb downstream of the *yihN* protospacer and about 30 kB upstream, 3 hours after *Ec*-IE induction. In contrast, genomic DNA remained intact in the AcrIE9-expressing cells (**Figure 7C, Supplementary Figure 6**). The results confirm that AcrIE9 blocks target DNA degradation *in vivo* and demonstrate how this Acr can be used to regulate chromosomal deletions by Type IE effector complexes.

### AcrIE9 and AcrIE10 form a widespread functional module co-occurring with a novel HTH-like protein

*Ec*_AcrIE9 and *Pa*_AcrIE9 are both highly active against the *E. coli* Type IE CRISPR-Cas system. A homolog of AcrIE9, named AcrIE9.2, was also recently described in *K. pneumonia*^37^. This protein demonstrates activity against *K. pneumonia* CRISPR-Cas of the subtype IE*, that is distinct from *Ec*-IE system. Compared to the *Kp*_AcrIE9.2, *Ec*_AcrIE9 contains a single amino-acid change, while it shares only ∼35% sequence identity with *Pa*_AcrIE9, which lacks a C-terminal extension (**Supplementary Figure 7A**). The corresponding sequences of the targeted CRISPR-Cas systems are also quite divergent, and Cas7 of *E. coli* shares only 32% and 27% amino-acid identity with Cas7 from *P. aeruginosa* and *K. pneumonia* (**Supplementary Figure 7B**). This suggests that despite high sequence divergence, AcrIE9 retained the ability to target a conserved interface within the Cas7 subunit of multiple Type IE systems, a feature that could provide a significant evolutionary advantage.

Indeed, AcrIE9 is the most widespread Acr protein among *Enterobacterales* (**Figure 1**), and we decided to further investigate its distribution and local genomic context. We extracted all *Enterobacterales* loci encoding AcrIE9 and classified them to groups based on the content of encoded proteins (see Methods, **Supplementary Figure 8**). AcrIE9 was most frequently found encoded on plasmids and conjugative plasmids (12,693 groups), and rarely in prophages and phages (29 groups), while chromosomal occurrences (1,023 groups) may also represent unrecognized MGEs (**Figure 8A**). Conjugative plasmids are known to encode hotspots of antiimmune proteins, including Acr clusters, that are expressed during transfer of ssDNA into the recipient cell^32,83^. While we were unable to predict ssDNA-specific promoters upstream of AcrIE9 operons, the location of AcrIE9 genes might support its role in the early suppression of the CRISPR-Cas response during plasmid conjugation. AcrIE9 is most prevalent in *Klebsiella*, followed by *Escherichia* and *Serratia* (**Figure 8B**), the structures of representative loci encoding AcrIE9 are provided in **Figure 8C**.

**Figure 8:**
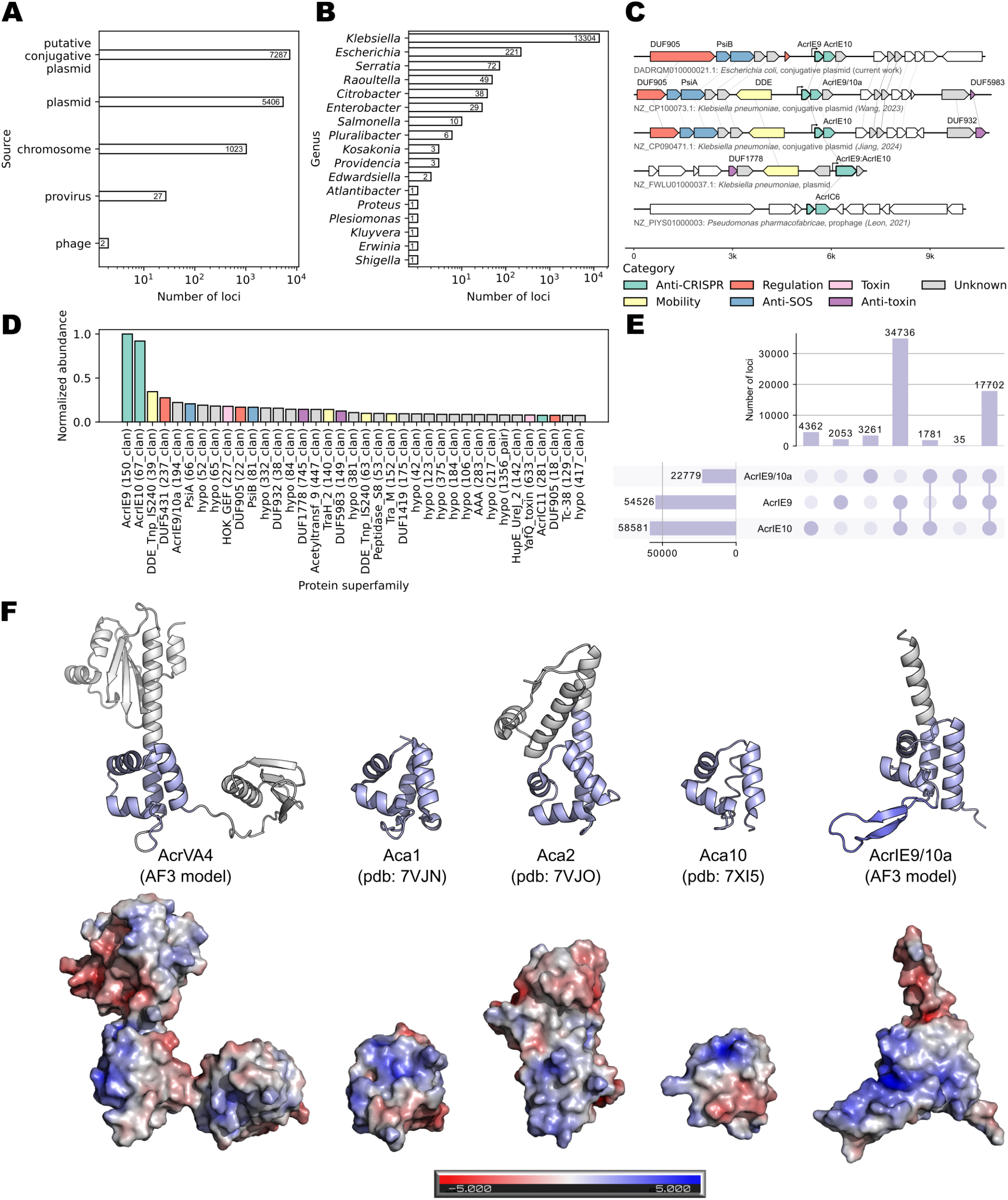
AcrIE9 and AcrIE10 form a highly abundant genetic module associated with a novel HTH-like protein. **(A-B)** Distribution of AcrIE9 across mobile genetic elements **(A)** and genera **(B)** within *Enterobacterales* order. **(C)** Representative genomic loci harboring *acrIE9*. Genes encoding proteins from one superfamily are connected with lines. σ70 promoters predicted with BPROM are shown as arrows. **(D)** Normalized abundance of most common protein superfamilies found in AcrIE9-encoding loci. Bars are colored according to manually assigned functional categories, identical to panel **(C). (E)** Co-localization of *acrIE9, acrIE10*, and *acrIE9/10a* in genomes of *Enterobacterales*. Genomic distance lower than 2 kB was accounted as co-localization. **(F)** Structure and surface charge distribution of AcrIE9/10a in comparison with distantly related Aca proteins and AcrVA4. Common HTH protein fold is colored in light blue. Comparison of AcrIE9/10a protein models with other Aca types is provided in **Supplementary Figure 9**. All used Aca protein sequences are listed in **Supplementary Table 3**.

We further investigated AcrIE9 potential functional partners based on their co-localization frequency (**Figure 8D**). The AcrIE9 was often encoded close to FlmC (DUF5431) and IS*Kpn28* DDE transposase^84^, suggesting the role of this element in the Acr mobility. However, the most widespread AcrIE9 partner is AcrIE10, recently described as a novel inhibitor of the *K. pneumonia* Type IE* CRISPR-Cas system that binds to the free Cas7* subunit^38^. These proteins were almost exclusively co-encoded together in our dataset, sometimes even forming a fusion (**Figure 8C**,**D**). AcrIE9 co-localizes with AcrIE10 in ∼96% of loci, and in ∼32.5% of instances this pair is flanked by a novel gene, encoding a protein of unknown function that we tentatively named AcrIE9/10-associated or AcrIE9/10a (**Figure 8E**). HHPRED analysis identified distant similarity of this protein with multiple HTH anti-CRISPR associated (Aca) transcriptional repressors: Aca1^85^, Aca2^85,86^, Aca10^87^, and AcrVA4^88,89^ protein (**Figure 8F**).

Since Acr operons are frequently controlled by Aca proteins, we questioned whether AcrIE9/10 expression is regulated. Given its genetic location and distant similarity with known Acas, we initially considered AcrIE9/10a as a transcriptional repressor. We modeled its structure with AlphaFold3 (AF3)^90^ and aligned it with known or predicted structures of all confirmed Aca proteins and AcrVA4 (**Figure 8F, Supplementary Figure 9A, Supplementary Table 3**). AcrIE9/10a is predicted to possess a helix-turn-helix motif with insertion of antiparallel beta-sheet, reminiscent of a winged HTH domains. Amino-acid conservation analysis confirmed the presence of beta-sheet inserts in all AcrIE9/10a homologs, while in a few instances, protein terminated in this region (**Supplementary Figure 9B**,**C**).

We cloned and produced AcrIE9/10a protein from a clinically-relevant *K. pneumonia* ST15 plasmid RJKP41^37^, that is nearly identical to *E*.*coli* AcrIE9/10a (**Supplementary Figure 7C**,**D)**, and performed a direct EMSA experiment with 5-FAM-labeled dsDNA substrate containing cognate *acrIE9* promoter. AcrIE9/10a demonstrated weak DNA binding that was completely prevented by an excess of non-labeled competing DNA (**Supplementary Figure 10**). Thus, although HTH-like AcrIE9/10a protein demonstrates non-specific DNA binding, it is unlikely to play a role of the Aca repressor for the AcrIE9/10 operon.

Our analysis establishes AcrIE9/AcrIE10 as the most widespread anti-CRISPR locus in sequenced *Enterobacterales* genomes and reveals an unusual co-occurrence of two Acr proteins targeting the same subunit of IE Cascade complex. In addition, we identified a novel protein with HTH-like architecture as a frequent component of the AcrIE9/AcrIE10 locus. Further research is needed to establish the role for each component of this locus and determine details of its transcriptional regulation.

## Discussion

In this study, we aimed to identify an anti-CRISPR protein active against the model Type IE CRISPR-Cas system of *Escherichia coli* K12. The biological role of this system remains enigmatic due to H-NS-mediated silencing of *cas* promoters and the absence of known conditions leading to activation of interference or adaptation^47,48^. However, the natural diversity of *E. coli* Type IE spacers suggests potential involvement in conflict^60^, which would imply that MGEs infecting *E. coli* could encode Acr proteins promoting their coexistence with the resident CRISPR-Cas systems. To test this hypothesis, we conducted a large-scale analysis of known Acr protein distribution within the *Enterobacterales* order, revealing an abundance of potential inhibitors targeting IB, IC, IE, IF, IIA, and VA CRISPR-Cas systems (**Figure 1B**). This finding aligns with the fact that many *Enterobacterales* species (e.g., *Klebsiella, Serratia, Pectobacterium*) serve as model organisms in studies of naturally active CRISPR-Cas systems^91,92^. Among Type IE inhibitors, AcrIE9 and AcrIE10 emerged as the most prevalent. We experimentally evaluated 11 representative AcrIE and AcrIF homologs against *E. coli* K12 Type IE system and demonstrated that AcrIE9 exhibits robust anti-CRISPR activity.

Originally identified as a prophage-encoded inhibitor of *Pseudomonas aeruginosa* Type IE CRISPR-Cas system^23^, AcrIE9 was recently implicated in the dissemination of *bla*_*KPC*_ carbapenem resistance plasmids in *Klebsiella pneumoniae* ST15^37^. This homolog (designated AcrIE9.2) shares ∼35% amino acid sequence identity with the *P. aeruginosa* variant and inhibits the Type IE* CRISPR-Cas system. *E. coli* AcrIE9 discovered in this work is nearly identical to AcrIE9.2 and is also encoded on conjugative plasmid, that are likely transferred between *Klebsiella* and *Escherichia*. Remarkably, despite low sequence conservation between the Cas proteins of *P. aeruginosa, E. coli*, and *K. pneumoniae* (**Figure S7**), *Pa*_AcrIE9 and *Ec*_AcrIE9 effectively inhibit diverse systems. This suggests that Acr proteins often evolve to target the most conserved interfaces of CRISPR-Cas complexes, in line with the evidence that non-homologous Type IE and IF Acrs, repeatedly target the same surfaces of the Cascade complex^15^.

Does the discovery of AcrIE9 as a potent inhibitor of *E. coli* K12 Type IE CRISPR-Cas support the biological role of this system in anti-MGE immunity? Although AcrIE9 is found in 726 *E. coli* genomes, its prevalence in *Klebsiella* (>52,000 genomes, **Extended Data File 1**) suggests stronger selection pressure from *Klebsiella* CRISPR-Cas systems, while *E. coli* could serve only as a temporary reservoir of AcrIE9-encoding conjugative plasmids. Otherwise, it can be speculated that, unlike the model K12 strain, other strains of *E. coli* could encode active Type IE CRISPR-Cas systems that drive the accumulation of Acr proteins in mobile elements.

Characterization of the anti-CRISPR mechanism revealed that AcrIE9 binds the Cas7 subunit of the assembled Cascade complex, disrupting target DNA recognition both *in vivo* and *in vitro*. Intriguingly, AcrIE9 itself exhibits DNA-binding activity, though this does not confer nonspecific protection against other nucleases, such as Cas9. While it remains unclear whether DNA binding activity contributes to the anti-CRISPR function, it can explain the moderate toxicity of the protein. Other Acrs, like AcrIF9, can promote Cascade complexes binding to non-specific DNA^93,94^. Our results indicate decreased affinity of the *Ec*Cascade complexes to the target DNA, yet we cannot exclude formation of the triple Cascade:AcrIE9:DNA complexes with non-target DNA.

We have shown that AcrIE9 is sufficient to inactivate Cascade complexes *in vivo*. However, AcrIE9 is almost invariably co-encoded with AcrIE10, another Type IE inhibitor recently reported to target *K. pneumoniae* Type IE and IE* CRISPR-Cas systems^38^. AcrIE10 binds the free Cas7 subunit to prevent Cascade assembly, and it is not clear if this mechanism provides protection from pre-existing surveillance complexes during the entry of conjugative plasmid to the host cell. The co-occurrence of two Acrs targeting the same subunit implies a novel strategy for CRISPR-Cas suppression, warranting further study of this widespread anti-CRISPR locus.

We hypothesize that two Acrs may function at distinct stages of plasmid conjugation: AcrIE9 could neutralize preassembled complexes at the onset of DNA transfer, while AcrIE10 could prevent the assembly of novel Cascade complexes at later stages. The clustering of multiple Acrs and Racrs, targeting the same CRISPR system, in other mobile elements suggests that coordinated suppression with multiple inhibitors may represent a widespread anti-CRISPR strategy^18,19^. Alternatively, AcrIE9 and AcrIE10 can act synergistically to achieve complete inhibition of CRISPR interference at the early stage of conjugation.

Stable coexistence of MGEs and CRISPR-Cas systems necessitates tight regulation to balance the negative costs of toxic Acr production against the risk of CRISPR-mediated cleavage^36^. Typically, Acr transcription is controlled by cognate Acr-associated (Aca) helix-turn-helix (HTH) transcription factors, and 10 Aca classes have been experimentally validated^95^. We identified an HTH-like protein, AcrIE9/10a, in one third of AcrIE9/10 loci, though it lacks sequence-specific binding affinity for *acrIE9* promoter and its function remains to be shown.

In summary, we analyzed anti-CRISPR proteins distribution in *Enterobacterales*, identified AcrIE9 as a potent inhibitor of *E. coli* K12 Type IE CRISPR-Cas system, and elucidated its mechanism of action via Cas7 binding and inhibition of the surveillance complex interaction with target DNA. Our study highlights that AcrIE9 is genetically linked with AcrIE10, thus forming the most prevalent anti-CRISPR locus in *Enterobacterales* MGEs. While the individual components of this operon have been studied^37,38^, future work should elucidate their collective roles under native expression conditions in conjugative plasmids. This task is particularly urgent given the role of AcrIE9/10 loci in the dissemination of antimicrobial resistance genes in pathogenic bacteria^37,38^.

## Materials and Methods

### Bacterial strains and phages

All strains of bacteria, plasmids, and phages used in this study are listed in **Supplementary Table 4**. For the initial test of anti-CRISPR activity, the KD263 strain^66^ and its derivatives were used. KD263 contains a g8 spacer targeting a protospacer in the M13 phage genome. The *cas3* gene expression is driven by the *lac* promoter, the remaining *cas* genes are transcribed from the *araBAD* promoter. SS80 is a derivative of KD263, with a spacer targeting T7 phage^44^. KD454 is a derivative of KD263 carrying a deletion of the *cas3* gene. Self-targeting experiments were performed in KD504, a derivative of KD263 with spacer targeting *yihN* gene. For testing Acrs in a natively active *Ec*-IE system, a pair of BW25113 derivative strains was used. BW39651 is a BW25113-derivative encoding T3 spacer targeting phage lambda (*λ*), while BW39671 is the same strain with the deletion of *hns* repressor gene^45^. Acr activity against the Ec_IF system was initially investigated in the LF82 *E. coli* strain^67^. Plasmid-encoded Ec_IF system was investigated in a CRISPR-deficient *E. coli* BB101 strain, susceptible to Mu phage infection^68^.

Bacterial cells were grown in LB (1% Bacto tryptone; 0.5% Bacto Yeast Extract; 1% NaCl) medium at 37°C overnight. Long-term storage stocks were prepared in LB supplemented with 10% glycerol and stored at -70 °C. To obtain a phage stock, an overnight bacterial culture was diluted 100-fold in LB medium and incubated at 37°C until OD600 reached 0.6. The culture was infected with phage at low MOI and further incubated overnight at 37°C. For phage *λ* production, the culture was supplemented with 0.2% maltose and 10 mM MgCl2. Phage M13 was produced on F’ KD263 host. Phage Mu was produced from a lysogenic culture, upon switching from 30°C to 45°C for 45 minutes at OD_600_=0.6. After induction, cells were grown at 37°C until lysis. The resulting lysates were purified from cell debris by centrifugation (6000g, 10 min, 4°C) and treated with chloroform (10 μl per 1 ml of lysate). Phage stocks were stored at +4°C.

### Plasmid construction

All primers used in this study are listed in **Supplementary Table 5**. Candidate anti-CRISPR genes were synthesized and cloned in a pBAD vector by Proteogenix company (France). Strep-tag II coding sequence with GSG linker (GSGSHPQFEK) was fused to the C-terminus of *Ec*_AcrIE9 and Pa_AcrIE9, encoded on pBAD via Gibson assembly (Gibson Assembly® Master Mix, NEB). Plasmid pFRG8 for Cascade-mediated sfGFP transcription silencing was constructed by KLD mutagenesis with the use of KLD reaction mix (NEB). Similarly, plasmid pFRC1T, promoting primed adaptation, was constructed from pFRG8 using KLD reaction. To enhance Acr expression, a variant of the pBAD_Ec_AcrIE9/Pa vector with modified RBS site was constructed via KLD reaction. These vectors were used in the pull-down experiments. For Ec_IF system expression, plasmid pET21d-LF82IFCR, kindly provided by A. Davidson, was modified to make it compatible with the pBAD vector. The ampicillin resistance gene was substituted with the streptomycin resistance gene via Gibson assembly (Gibson Assembly® Master Mix, NEB).

For heterologous expression and preparative purification of *Pa*_AcrIE9 protein, corresponding *acr* gene was re-cloned in pET28 vector under control of T7 promoter and in frame with C-terminal 6His-Tag sequence. *Ec*_AcrIE9 gene was re-cloned from plasmid pBAD-AcrIE9_Ec using primers 5’- GGAATTAATCATGAACTTCACCATTAAGAGTCGC-3’ and 5’-GCAGATCTCGAG-TTACTGATTATCTTCGC-3’ into the expression vector pET30 restricted by NcoI and XhoI enzymes. AcrIE9/10a from RJKP41 plasmid was cloned with primers 5’-CATGCCATGGCCATGAGTGACGTACTTTCCTG-3’ and 5’-CCGCTCGAG- TTACTTATCATTGTTTTCACTCACATTATTCGTTTCAAACGAGACAAG-3’ into pET28a-SUMO2 cloning vector restricted by NcoI and XhoI enzymes. The target plasmids pJ3 and pλ1-on-target were generated for gel shift binding assay with J3-Cascade and cleavage activity assay for SpCas9 nuclease, respectively. The pT7Blue-based plasmid pG8_dir carrying a 209-bp M13 fragment^69^ was used to replace g8 protospacer sequence 5′- GCTGTCTTTCGCTGCTGAGGGTGACGATCCCGC-3’ with J3 protospacer 5’- GCCAGTGATAAGTGGAATGCCATGTGGGCTGTC-3’ resulting in pJ3 plasmid^48^. The plasmid pλ1-on-target was obtained by blunt-end cloning synthetic DNA containing Cas9 protospacer^96^ in pT7Blue vector. For preparation of J3-Cascade (Cascade loaded with crRNA containing J3 spacer) we constructed pCDF- casBCDE(J3) plasmid by replacing g8 spacer in pCDF-casBCDE(g8)^97^ with J3 spacer^48^.

All constructs were transformed into XL1 Blue (Evrogen, Russia) or NEB® 10-beta (NEB) commercial chemically competent *E. coli* cells. Sequences of assembled plasmids were confirmed by Sanger sequencing (Genotech, Russia).

### Efficiency of plating (EOP) assay

To estimate the level of CRISPR-Cas defense and Acr activity, the titer of phage particles was measured by the double agar plating method. Plates were prepared with two layers: the bottom layer contained 1,5% LB agar media, while the top layer was mixed with overnight bacterial cell culture and included corresponding antibiotics and inductors, if necessary (1 mM IPTG, 0,2% Arabinose). For phage λ infection, the top layer of agar also contained 0.2% maltose and 10 mM MgCl_2_; for phage Mu infection, the top agar was supplemented with 10 mM MgCl_2_. Phage lysate was spotted on agar top in serial ten-fold dilutions. Plates were incubated overnight at 37°C. EOP with BW39651 and BW39671 strains were carried at 30°C. EOP with the phage T7 were carried at room temperature.

### Monitoring dynamics of phage infection in liquid culture

The overnight liquid cultures were diluted 100-fold and grew in liquid LB media at 37°C until OD_600_ = 0.6 with required antibiotics and in the presence/absence of inductors. The cultures were equilibrated to OD_600_ = 0.3 and 200 μl of equilibrated culture was loaded to 96 well plates and infected with the phage at indicated MOI (Multiplicity of Infection). The plate was loaded to the EnSpire Multimode Plate Reader (PerkinElmer, USA). Optical density was monitored for 20 hours at 37°C with intense orbital shaking (200 rpm).

### Phage release

To measure phage burst and accumulation of phage particles in liquid media, an extended one-step growth curve assay was performed. The overnight cultures of *E. coli* BW39671 encoding *λ*T3 spacer (and BW39651 as control) containing pBAD plasmids encoding Acrs were diluted 100-fold in 5 ml in liquid LB media supplemented with corresponding antibiotics before OD_600_ reaches 0.6. The cultures were mixed with phage *λ*_*vir*_ to achieve MOI=10^-3^. Aliquots were collected immediately after phage addition, and 1, 3, 5 hours post-infection. 600 μl of liquid media was collected, centrifuged at 13.000g for 5 minutes to remove cell debris and supernatant was plated in serial ten-fold dilutions on a lawn sensitive BW25113 *E. coli* strain to enumerate phage titer. The experiment was performed in three replicates.

### Self-toxicity inhibition assay

The overnight culture of *E. coli* KD504 cells was diluted 100-fold and grew in liquid LB media supplemented with corresponding antibiotics before OD_600_ reaches 0,2-0,3. At this point, 0,2% L-Arabinose and 1 mM IPTG were added to induce CRISPR and Acr expression. At every chosen time point 100 μl of cultures were collected from each sample and plated in a serial ten-fold dilutions on plates with 1% agar M9 media. M9 was prepared by diluting autoclaved 5x M9 salts (250mM Na_2_HPO_4_, 100mM KH_2_PO_4_, 50mM NaCl, 10mM NH_4_Cl) in mQ and mixing with MgSO_4_ (2 mM final concentration) and CaCl_2_ (0.1 mM final concentration). This media was additionally supplemented with 1% agar and 0,2% D-Glucose as carbon source. Plates were incubated at 37°C overnight. For NGS sequencing, 5 mL of cell culture aliquots were collected to extract genomic DNA. Genomic DNA purification was performed with Genomic DNA Purification Kit (Evrogen) under manufacturer protocol.

### Naive adaptation assay

*E. coli* BL21AI culture was co-transformed with plasmids pCDFCas1-2 and pBAD empty vector or pBAD_Pa_AcrIE9/Ec by electroporation. The next day, a single colony from each plate was inoculated into liquid LB media with corresponding antibiotics and grew until OD_600_ = 0,2-0,3. Cells were pelleted by centrifugation (3.200 g at room temperature) and put in 5 ml of fresh LB media without antibiotics. Cas1-2 and Acr expression was induced with 1 mM IPTG and 0,2% L-arabinose and following induction 100μl cell aliquots were collected each hour by centrifugation at (3.200 g at 4°C) and stored at -80°C in 10% glycerol. The naive adaptation was detected by PCR amplification from 100-fold diluted bacterial culture, using leader - repeat pair of primers Moji3 and Moji4 (**Figure 4A, Supplementary Table 5**). PCR products were loaded on 2% TAE Agarose gel and ran for 1 hour at 70W.

### Primed adaptation assay

*E. coli* KD263 culture was co-transformed with pFR66_G8(C1T) and pBAD empty vector or pBAD_Pa_AcrIE9/Ec by electroporation. The next day, a single colony from each plate was inoculated into liquid LB media with corresponding antibiotics and grew overnight at 37°C. Overnight culture was diluted 100-fold and grown in a fresh LB media without antibiotics until OD_600_ reaches 0.2-0.3. Then, 0,2% L-arabinose and 1 mM IPTG were added to induce CRISPR-Cas and Acr expression. 3h and 24h after induction, cell aliquots were collected by centrifugation at (3.200 g at 4°C) and frozen at -80°C in 10% glycerol. The primed adaptation was detected from 100-fold diluted bacterial culture, by PCR amplification using leader - spacer (G8) pair of primers EcLdr_F and M13G8_R (**Figure 4B, Supplementary Table 5**). PCR products were loaded into 2% TAE Agarose gel and ran for 1 hour at 70W.

### sfGFP transcriptional silencing

To investigate *Ec*-IE Cascade DNA binding activity *in vivo*, we measured sfGFP expression from a promoter overlapping with *Ec*-IE protospacer. *E*.*coli* KD263 strain with G8 spacer carried pBAD empty vector or pBAD encoding Pa_AcrIE9/Ec, and sfGFP-encoding plasmid with (pFR66) or without (pFR66) a G8 protospacer. The overnight liquid cultures were diluted 100-fold and grew in LB supplemented with required antibiotics/inducers at 37°C until OD_600_ reaches 0.6. The cultures were equilibrated to OD_600_ = 0.3 and 200 μl was loaded to 96 well plate. Cell growth continued in EnSpire Multimode Plate Reader (PerkinElmer, USA) at 37°C with monitoring sfGFP fluorescence (Ex λ = 480, Em λ = 510) and absorbance (OD_600_). The resulting fluorescence data was normalized to an optical density. The experiment was performed in three replicates.

### Protein pull-down

Overnight culture of *E. coli* BW25113 carrying pBAD_Pa_AcrIE9/Ec C-Strep was diluted 100-fold in 3 L of LB media with corresponding antibiotics. When the culture reached OD_600_ = 0.2-0.3 CRISPR-Cas and C-terminally Strep-tagged AcrIE9 were induced with 1 mM IPTG and 0,2% L-Arabinose were added and cells grew overnight at 37°C. Thereafter, cells were spun down by centrifugation at 3500 g for 20 min at 4°C and cell pellets were resuspended in StrepA buffer (20 mM Tris-HCl pH 8.0, 50 mM NaCl, 5 mM βmercaptoethanol, 1 mM EDTA) supplemented with 5 mg/ml of lysozyme and protease inhibitor cocktail (Roche). Cells were lysed by sonication on ice for 30 min with using 10s impulse - 20s rest circle at power 20% (Qsonica sonicator with 6 mm sonotrode). Cell debris was removed by centrifugation at 25.000 g for 45 min at 4°C. Purification of Strep-tagged protein together with suggestive partners was carried out with a BioRad NGC Chromatography System on a 5 mL StrepTrap HP column (GE Healthcare). Before applying the bacterial lysate, the column was equilibrated with a StrepA buffer. Proteins bound to the column were eluted with a StrepB buffer (StrepA supplemented with 2.5 mM desthiobiotin). The protein composition of the eluted fractions was analyzed by Laemli SDS-PAGE after concentration with 3-kDa ultra centrifugal Amicon filters. The identity of protein bands was determined by matrix-assisted laser desorption/ionization time-of-flight (MALDI-TOF) mass spectrometry. Samples were treated with Trypsin Gold (Promega) in accordance with manufacturer’s instructions. Mass spectra were obtained using the rapifleX system (Bruker).

To purify Ec_IE Cascade from cell culture expressing untagged *Pa*_AcrIE9, *E*.*coli* BL-21 AI cells were transformed with plasmids pWUR515^98^ (Strep- CasC,CasD,CasE), pWUR407, and EcCRISPR P7-7x, and pET30-AcrIE9_Pa, and plated on LB agar supplemented with appropriate antibiotics. Expression of *Pa*_AcrIE9, Cascade and CRISPR RNA was induced using 0.2% Arabinose and 0.5mM IPTG when cells reached an OD_600_ of 0.4. After expressing at 37 °C for 16 hours, cells were harvested by centrifugation at 5,000G for 20 minutes. Cell pellets were resuspended in lysis buffer containing 20 mM Tris (pH 8.0), 100 mM NaCl, and 1 mM DTT before lysing by sonication. Cellular debris was removed by centrifugation at 26,892g for 30 min and the cleared lysate was loaded onto a 5mL StrepTrap HP column. After washing with 15 column volumes of lysis buffer, the Cascade complex was eluted using lysis buffer supplemented with 50mM desthiobiotin. Elutions were concentrated before being resolved on a Superose 6 10/300 size exclusion column equilibrated in SEC buffer (lysis buffer + 5% glycerol). Fractions from the main peak were concentrated to ≈6uM and proteins were visualized on a 12% acrylamide gel via SDS-PAGE.

### Protein purification

Cascade (Cas8,11,7,5,6) or Cascade subcomplexes lacking Cas8 (Cas11,7,5,6), or Cas8 and Cas6 (Cas11,7,5) were prepared from E. coli KD418 cells co-expressing appropriate *cas* genes with a source of crRNA (**Table S4**) and were one-step affinity-purified on Strep-Tactin® column (IBA) using the N-terminal Strep-tag attached to the Cas11 subunit^99^ according to manufacturer’s protocol.

N-terminal 6His-tagged Cas8 protein was IMAC purified from *E*.*coli* BL21 Star (DE3) pLysS strain (Invitrogen) transformed with pET30-*casA* expression plasmid (**Supplementary Table 4**) followed by gel-filtration on Superose-6 column (Pharmacia) equilibrated with 40 mM Tris-HCl buffer (pH8) containing 200 mM NaCl, 5% Glycerol, 1mM TCEP.

C-terminal 6His-tagged *Pa*_AcrIE9 protein was IMAC purified from *E*.*coli* Rosetta 2 (DE3) strain (Novagen) transformed with pET28-*Pa*_AcrIE9 expression plasmid (**Supplementary Table 4**) followed by ultrafiltration through VivaSpin 5 kDa MWCO concentrator (EMD Millipore) for concentrating and buffer exchange in 40mM Tris-HCl, pH 7.5, 300 mM NaCl, 1 mM TCEP, 5% Glycerol.

N-terminal 6His-tagged AcrIE9_Ec protein was IMAC purified from *E*.*coli* Rosetta 2 (DE3) strain (Novagen) transformed with pET30-AcrIE9_Ec expression plasmid (**Supplementary Table 4**) followed by ultrafiltration through VivaSpin 5 kDa MWCO concentrator (EMD Millipore) for concentrating and buffer exchange in 40mM Tris-HCl, pH 7.5, 300 mM NaCl, 1 mM TCEP, 5% Glycerol. Before use in the binding reactions with target plasmid DNA, the N-terminal 43 aa tag has been removed by Enterokinase, light chain (NEB) according to the manufacture protocol.

N-terminal 6His-SUMO-tagged AcrIE9/10a protein was IMAC purified from *E*.*coli* BL21 (DE3) strain (Novagen) transformed with pET28a-AcrIE9/10a expression plasmid (**Supplementary Table 4**) followed by ultrafiltration through VivaSpin 5 kDa MWCO concentrator (EMD Millipore) for concentrating and buffer exchange in 40mM Tris-HCl, pH 7.5, 300 mM NaCl, 1 mM TCEP, 5% Glycerol. Before use in the binding reactions with target DNA, His6-SUMO tag was removed by SENP1 protease digestion overnight followed by Ni-NTA purification to obtain tag-free AcrIE9/10a.

SpCas9 nuclease from *Streptococcus pyogenes* was expressed in *E*.*coli* Rosetta 2 (DE3) strain transformed with pMJ806 (Addgene) and purified by protocol published earlier^100^. To obtain catalytically active Cas9 nuclease we used synthetic guide RNAs: tracrRNA and crRNA ordered in IDT (**Supplementary Table 5**). Reconstituted complex SpCas9/tracrRNA/crRNA was validated for binding and cleavage activity towards a target plasmid DNA.

### Glutaraldehyde crosslinking between AcrIE9 and E.coli IE Cascade

Crosslinking between *Pa*_AcrIE9 and Cascade or its sub-complexes was initialized by addition of glutaraldehyde (GA) to a final concentration of 0.5% into the mix of 20 μM *Pa*_AcrIE9 and 1 μM Cascade in the buffer containing 40 mM MOPS, pH 8.0, 50 mM NaCl at room temperature for 2-3 min. Then reaction was quenched by addition of TRIS solution to a final concentration of 100 mM. Crosslinking products were resolved by denaturing SDS-PAGE and analyzed by mass spectrometry.

### Electrophoretic Mobility Shift Assay (EMSA)

J3-Cascade was mixed with *Pa*_AcrIE9 or *Ec*_AcrIE9 in buffer containing 40 mM Tris-HCl (pH7.5), 50 mM NaCl, 0.5 mM TCEP, and 0.1 mg/ml BSA for 10 min at 37°C, then target plasmid DNA pJ3 was added and incubated for 20 min at 37°C. Samples containing Cascade:DNA complexes engaged by AcrIE9 were run in 0.7% TBE-agarose gel for 3 hours at 60V, 30 mA, 2W. Then the gel was stained with EtBr and visualized by GelDoc XR+(Bio-Rad).

Binding reaction between *Pa*_AcrIE9 and 52-bp short single-stranded and double-stranded oligos labeled by fluorescein (**Supplementary Table 5**) were carried out in the same binding buffer and samples were loaded on native precast 10 to 20% gradient polyacrylamide gel (Novex, Invitrogen), and run in trisglycine buffer at 20 mA for 2 hours. The bound complexes were visualized with GelDoc XR+(Bio-Rad).

Binding reaction between AcrIE9/10a and *acrIE9* promoter reaction was performed using 300 bp PCR product amplified with 5-FAM labeled forward primer (**Supplementary Table 5**). EMSAs were performed in a reaction volume of 20 μl containing binding buffer (5% glycerol,pH 7.8 20 mM Tris HCl, 150mM KCl, 2 mM MgCl_2_, 0.1μg/μl BSA,0.5 mM DTT), 20 nM of *acrIE9* promoter substrate and a gradient concentration of Aca14 (range 0–800 nM). 0.5μl of herring sperm DNA (5mg/ml, Sangon Biotech (Shanghai) Co., Ltd.), was used to exclude non-specific binding of the protein to DNA. Binding reactions were mixed on ice and incubated for 30 min. Then 16 μL of each reaction was loaded on a 5% Tris-Glycine native PAGE gel and run at 130 V for 55 min on ice. An Amersham TyphoonTM 5 Biomolecular Imager was used for gel analysis.

### SpCas9 nuclease cleavage activity assay

For cleavage experiments, 75 nM of SpCa9 and gRNA (tracrRNA+crRNA) were preincubated at room temperature for 10 min in binding buffer containing 40mM Tris-HCl(pH7.5), 100 mM NaCl, 1mM DTT, 5mM MgCl2, before initiating reaction by addition of 1.2 nM of target plasmid DNA pl1-on-target with or without 14 μM of AcrIE9 for indicated time at room temperature. Cleavage products were analyzed by 0.8% agarose gel electrophoresis and visualized by GelDoc XR+(Bio-Rad) after EtBr staining.

### Proteomic analysis

Protein-containing fractions obtained from pull-down with Strep-tagged AcrIE9 variants (or non-tagged proteins as controls) were loaded on 5% SDS-PAGE after boiling for 10 minutes in standard SDS-loading dye. Proteins were allowed to completely enter the resolving gel then the current was stopped. The first 3-4 mm of resolving gel containing the samples were sliced and digested with Trypsin Gold (Promega). To elute peptides, gel was incubated in a microtube shaker with 50 μl of 50% acetonitrile/5% formic acid for 45 min at room temperature. Liquid was removed and extraction was repeated. Final extraction step was carried with 50 μl of 90% acetonitrile/5% formic acid for 5 min. Collected peptides were vacuum dried and solubilized in 5 μl of 0.1% formic acid. Peptides were subjected to LC-MS/MS analysis with Q Exactive HF-X mass spectrometer (Thermo Scientific) as described before^101^. Raw data was processed with MaxQuant software and peptides were searched against *E*.*coli* BW25113 proteome extended with AcrIE9 sequences.

### NGS library preparation and sequencing

Sequencing libraries were constructed from 1 μg of total genomic DNA using MGI Easy PCR-Free Library Prep Set in accordance with User Manual v1.1 (MGI Tech Co., Shenzen, China), size-selected and cleaned up with the provided DNA Clean Beads. Quality control was performed with Agilent High Sensitivity D1000 ScreenTape Assay on 4200 TapeStation System (Agilent Technologies Inc., Santa Clara, CA, USA) and QuDye ssDNA kit (Lumiprobe, Moscow, Russia). Barcoded libraries were pooled and sequenced using DNBSEQ-G400 in 2x150bp PE mode.

### Self-targeting NGS data analysis

The obtained reads were trimmed and filtered with fastp v. 0.23.2^102^. Then reads were aligned on a reference genome (*Escherichia coli* KD403^103^ and cloning vector pBAD) with BWA mem v. 0.7.18^104^. Optical duplicates were marked with Picard v. 2.18.7. The alignments were sorted and filtered with samtools v.1.21^105^ to retain only primary aligned paired reads (‘-f 2 -q 20 -F 1024,256,2048’). The coverage was extracted for each position in bacterial genome and normalized with 1x normalization with bamCoverage utility from deepTools v.3.5.5^106^. Effective genome size was calculated with `uniquekmers.py` from khmer v.2.1.1^107^. Ratios of normalized coverage for samples with/without *Pa*_AcrIE9 expression in each time point were calculated with bigwigCompare from deepTools.

### Identification of known anti-CRISPRs within *Enterobacterales* order

#### Search for known anti-CRISPR proteins in non-redundant protein database

An exhaustive search of known Acr proteins in non-redundant protein database was performed. hmmsearch v.3.3.2 (http://eddylab.org/soft-ware/hmmer/) was used to scan non-redundant protein sequences of bacteria, viruses, and plasmids (taxid: 2, 10239, and 36549 from nr BLAST, v.5) against HMM profiles of Acr proteins from a database of anti-prokaryotic immune systems (dbAPIS) v.1^35^. The protein hits with E-value less than 1e-10 were considered as Acrs and were included in further analysis.

#### Taxonomic and genetic localization assignment for acr genes

For each obtained hit we explored contigs, in which they are encoded, and defined their taxonomy based on sequence metadata. Taxon ID for genome assemblies were obtained with NCBI datasets toolkit v.16.21.0^108^, and the corresponding lineages were fetched with taxonkit v. 0.14.2^109^. Contigs encoding potential Acrs of *Enterobacterales* were selected, and mobile genetic elements were predicted with geNomad v. 1.8 using end-to-end pipeline^110^.

#### Loci extraction

Genome assemblies of *Enterobacterales* contigs harboring Acrs were fetched with `datasets download` tool from NCBI datasets toolkit v.16.21.0. Contigs encoding Acrs were clustered with MMseqs2 v. 15-6f452 cluster^111^ to remove redundancy with following parameters: minimal coverage 0.9, minimal sequence identity 0.99, clustering mode 0, coverage mode 0. Then the nucleotide sequences of Acrs genomic contexts (5 kb upstream and downstream) were extracted and used for further analysis of Acrs genomic context.

### Analysis of AcrIE9 genomic context

#### Search for proteins sequence homology groups

Coding regions in extracted genomic loci containing Acr hits were predicted with prodigal v. 2.6.3^112^. Predicted protein sequences were clustered with MMSeqs2 v. 15-6f452^111^ with the following parameters: minimal sequence identity 0.3, minimal coverage 0.8, clustering mode 0, coverage mode 0. Set of experimentally validated Acr sequences were added to the clustering procedure to remove possible falsepositive hmmsearch hits. Next, multiple sequence alignments (set of MSAs) for each protein cluster were extracted, and MSAs for clusters with more than 3 representatives were aggregated into a HH-suite3 database v. 3.3.0^113^. Set of MSAs were searched against formed database of HMM-profiles resulting in a list of pairs of homologous clusters (E-value threshold 5e-5). Communities, or locally dense subgraphs, were obtained from this similarity network with ClusterONE v.1.2^114^, and were considered as protein clans (or protein superfamilies).

#### Loci diversity analysis

The diversity of local genomic contexts for AcrIE9 was described as follows. To leverage the effect of disproportionate abundance of different bacterial species sequences, groups of loci similarity were defined. First, the target loci were represented as sets of protein clans, encoded in these loci. Second, Jaccard similarity between each pair of loci was calculated. Third, for estimation of similarity threshold, Jaccard similarity for 200 permuted sets were calculated (protein clans were randomly permuted among loci). The value of 0.01 of permuted 99th quantiles observed in permutations was selected as a threshold (0.4), and all pairs of loci with Jaccard similarity more than the threshold were considered as edges within the loci content similarity network. Same as for protein cluster similarity network, communities of the network were found. For each of 67 found communities, the within-community abundance of protein clans was defined as the proportion of loci within the community, in which the genes corresponding for protein clans are present. Then the normalized abundance was calculated for each protein clan as a sum of within-community abundance divided by the number of communities. Given that the abundance of protein groups is calculated only for loci encoding AcrIE9 homologs, the higher normalized abundance reflects higher AcrIE9 co-localization rate with respect to a more variable 10 kbp genomic context. For the analysis of genomic context, only proteins with normalized abundance higher than 0.075 were selected.

#### Construction of HMM profiles

For the analysis of AcrIE9, AcrIE10 and AcrIE9/10a co-occurrence, we performed an HMM profile search against the nr BLAST database. To build HMM profiles, target protein sequences were extracted for each protein superfamily with seqkit v2.8.2^115^, and clustered with MMSeqs2 to remove redundancy with following parameters: minimal coverage 0.9, minimal sequence identity 0.95, clustering mode 0, coverage mode 0. Resulting non-redundant sequence sets were aligned with MAFFT v.7.490^116^, and then converted into HMM profiles with hmmbuild from HMMER package.

#### Protein sequences annotation

Alignments of protein clusters obtained with MMSeqs2 were searched against Pfam-A database^117^ with hhsearch tool from HH-Suite3 v.3.3.0^113^ (E-value threshold 1e-5). Then the hit with lowest E-value was assigned as a true hit for each protein cluster. Of note, such an approach excludes assignment of more than one found domain to the protein cluster.

#### AcrIE9 promoters’ prediction

Upstream regions of AcrIE9 gene for four representative loci were selected with bedtools getfasta v2.30.0^118^ (-200nt and +50nt with respect to gene start). The promoter regions within obtained nucleotide sequences were predicted with BRPROM^119^.

#### AcrIE9/10a identification and structure comparison analysis

Amino acid sequence of AcrIE9/10a protein was queried to the HHPRED WebServer^120^. After the homology was established, the protein structure was predicted with AlphaFold3^90^ (WP_013023810.1 was used as a representative AcrIE9/10a), and the obtained molecule model was searched with DALI^121^ against the PDB database. To evaluate whether AcrIE9/10a belongs to one of the previously described Aca classes, the protein structure comparison was performed. Protein structures of all known Acas were modeled with AlphaFold3 (Protein sequences/structures listed in **Supplementary Table 3**). To build **Figure 8F**, alignment of 3D structures was performed with jFATCAT^122^ in a rigid mode (all-vsAcrIE9/10a), while for the analysis presented at the **Figure S9A**, FoldMason WebServer^123^ in all-against-all mode was used. Resulted structural alignments were visualized with PyMOL v.3.0.0. The electrostatic potentials obtained with APBS^124^. For the coloring of protein structures according to LDDT for MSA, the custom python3 script was utilized, and available on GitHub (https://github.com/ombystoma-young/annotationtoolkit/foldma-son2pdb). The global alignment of homologs of AcrIE9/10a was performed with MAFFT on a subset of deduplicated sequences of AcrIE9/10a, belonging to protein clusters with a number of representatives more than 2 (resulted in alignment of 89 unique sequences).

### Workflow

We processed data and visualized results using Python3 v.3.11.2 (https://docs.python.org/3.11/) and R v.4.2.3 (https://www.R-project.org/). To work with networks, we used the NetworkX v.2.7.1^125^. The multiple sequence alignments were visualised with pyMSAviz v.0.4.2 (https://github.com/moshi4/pyMSAviz). We aggregated the resulting figures and drew a model schema with the help of Inkscape v.1.3.2 (https://inkscape.org/credits/) or Biorender. The proposed workflow is formalized in a Snakemake-based pipeline (Snakemake v.7.25.2^126^). The developed pipeline and all supplementary scripts are available at GitHub: https://github.com/ombystoma-young/loci-search-pipe.

## Acknowledgements

Mass-spectrometry was performed at Skoltech Advanced Mass-Spectrometry Core Facility with support of the internal Skoltech grant, we thank Dr. Maria Zavialova for the samples handling. We thank Dr. Alan Davidson, who kindly shared *E. coli* Type IF CRISPR-Cas system plasmids. We express deep gratitude to Dr. David Bikard and members of his laboratory for the support of this research project.

## Author contributions

Conceptualization, A.I.; Molecular cloning, microbiology and phage methods, D.T., S.F., D.Y.; Bioinformatic analysis, O.K., and V.M.; *In vitro* assays, K.K., K.V., D.L., and N.B.; BGI sequencing, A.D.; Resources, A.I., B.W., M.W., E.S., K.S.; Writing—original draft, A.I., D.T., O.K.; Writing—review and editing, A.I., B.W., E.S., K.S; visualization, D.T., O.K. All authors have read and agreed to the published version of the manuscript.

## Data availability statement

Plasmids and bacterial strains used in this work are available upon request from the lead contact, Dr. Artem Isaev.

---

**Figure S1.**
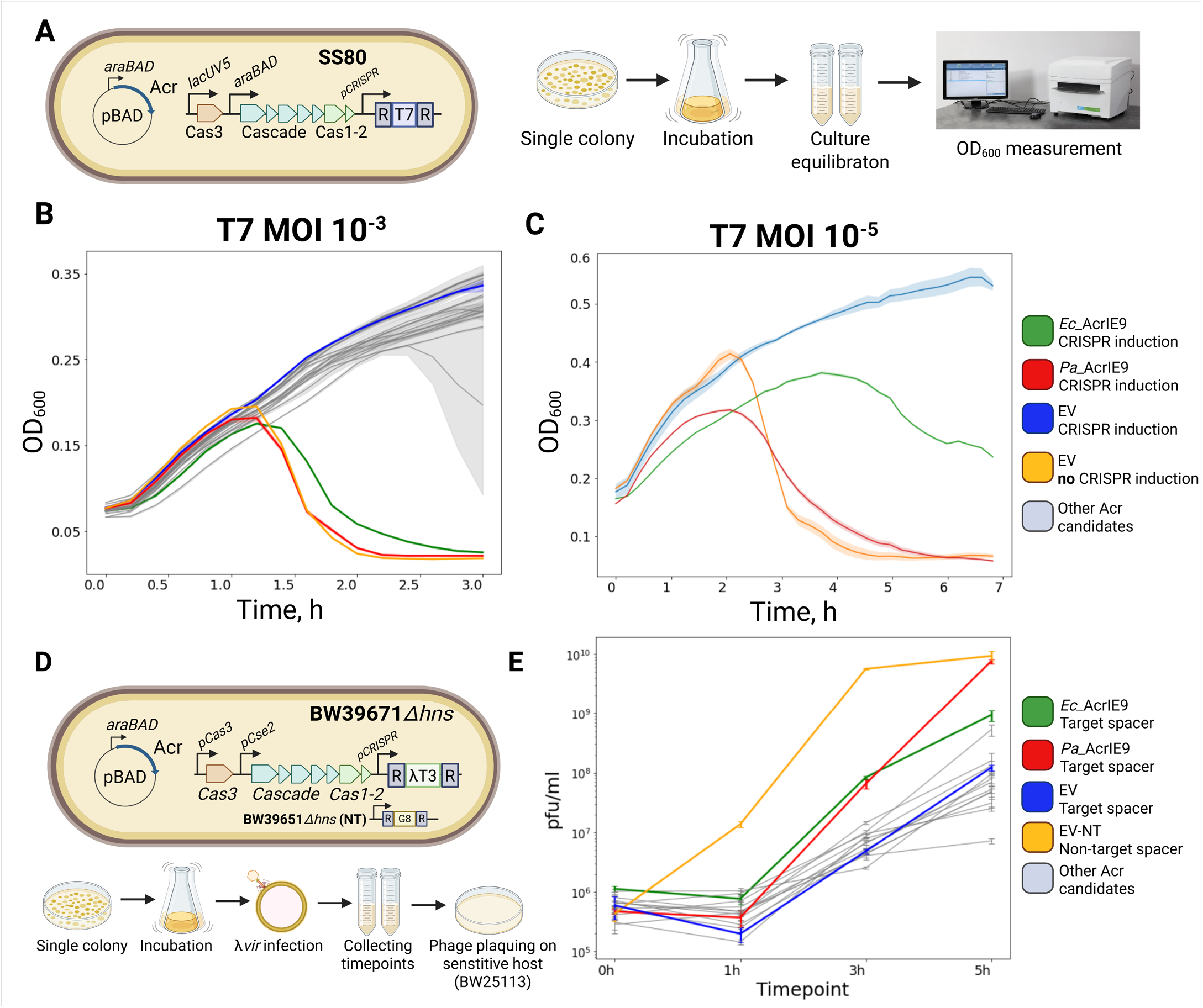
Experimental validation of *Pa*_AcrIE9 and *Ec*_AcrIE9 *Ec*-IE-inhibiting activity. **(A)** *E. coli* SS80 strain, a KD263 derivative with T7-targeting spacer, was used for T7 infection assay. **(B-C)** Growth curves of SS80 Culture infected with T7 in conditions of Acr expression. Cells were grown with induction until OD_600_ ∼0.6, equilibrated to OD_600_ = 0,3 and infected with the T7 phage at MOI=10^-3^ **(B)**, MOI=10^-5^ **(C). (D)** *E. coli* BW39671 *Δhns* strain, a BW25113 *Δhns* derivative with a *λ* phage-targeting spacer T3, was used for *λ*_*vir*_ phage infection assay. **(E)** Dynamics of phage *λ*_*vir*_ release from infected *E. coli* BW39671 *Δhns* cells in conditions of Acr expression. Cells were grown at 30°C with induction until OD_600_=0.6 and then infected with *λ*_*vir*_ at MOI=10^-2^. Phage titer in the spent media was measured at indicated time points on a lawn of sensitive *E*.*coli* BW25113 host. Acr expression was induced with 0.2% L-arabinose, CRISPR-Cas system was expressed from native promoters. All experiments were performed in three biological replicates.

**Figure S2.**
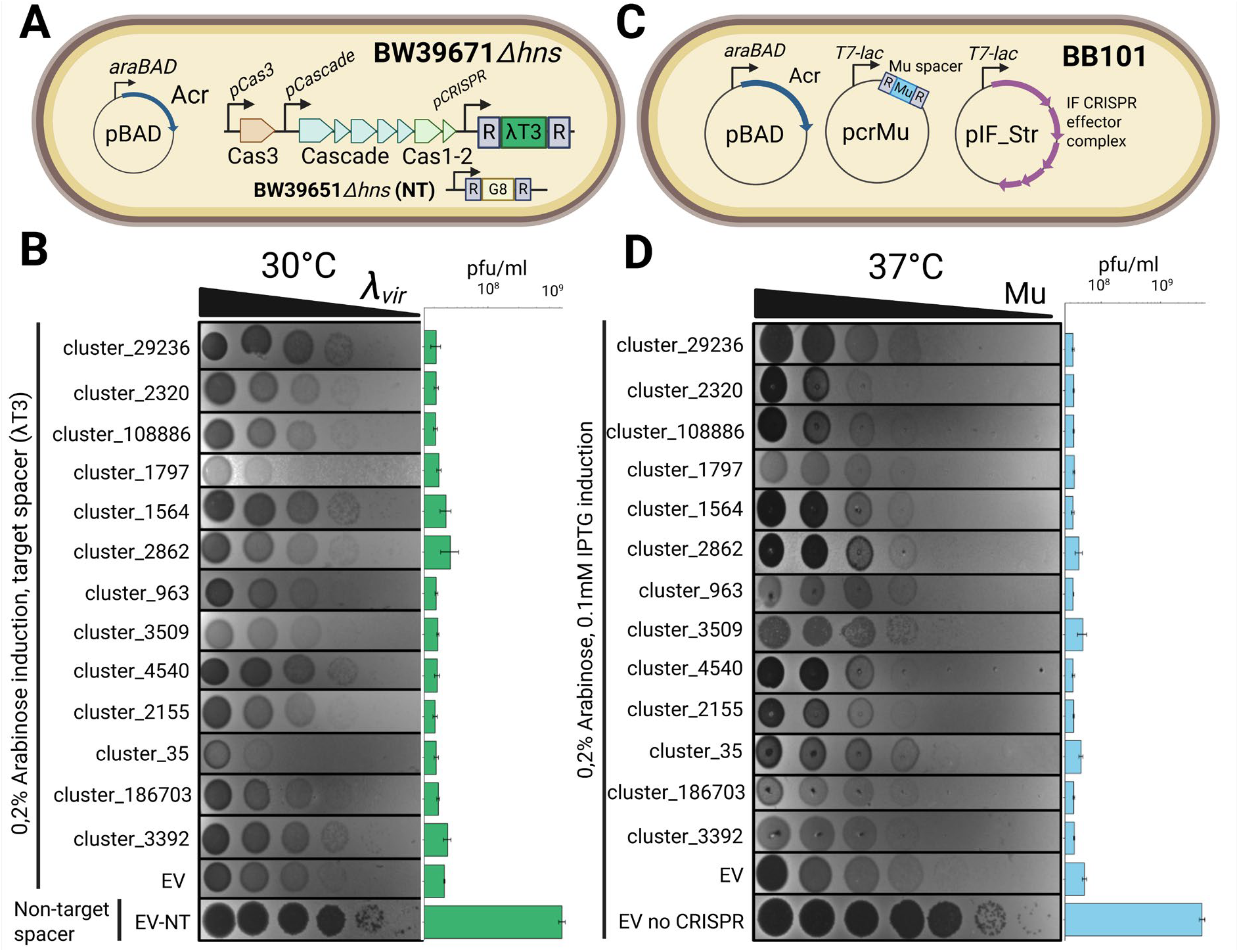
Screening of ML-predicted collection of anti-CRISPR candidates. **(A)** *E. coli* BW39671 *Δhns* strain, a BW25113 *Δhns* derivative with a *λ* phage-targeting spacer T3, was used for *λ*_*vir*_ phage infection assay. **(B)** EOP with phage *λ*_*vir*_ on the lawns of BW39671 *Δhns* cells with induced expression of Acr ML-predicted candidates. Cultures without a *λ*_*vir*_ targeting spacer (EV-NT) and with empty pBAD vector (EV +ind) were used as controls. **(C)** *E. coli* BB101 strain with a plasmid-encoded IF LF82 system and phage-targeting spacer was used for the phage Mu phage infection assay. Acr expression was induced with 0.2% L-arabinose, CRISPR-Cas expression was induced with 0.2% L-arabinose and 1 mM IPTG. **(D)** EOP with phage Mu on the lawns of BB101 cells with induced expression of Cas proteins and Acr ML-predicted candidates. Cultures with empty pBAD vector (EV-NT) and without CRISPR-Cas encoding plasmid (no CRISPR) were used as controls. Acr expression was induced with 0.2% L-arabinose, CRISPR-Cas and spacer expression were induced with 0.1 mM IPTG. All experiments were performed in biological triplicates and bars on the right represent mean pfu/ml values.

**Figure S3.**
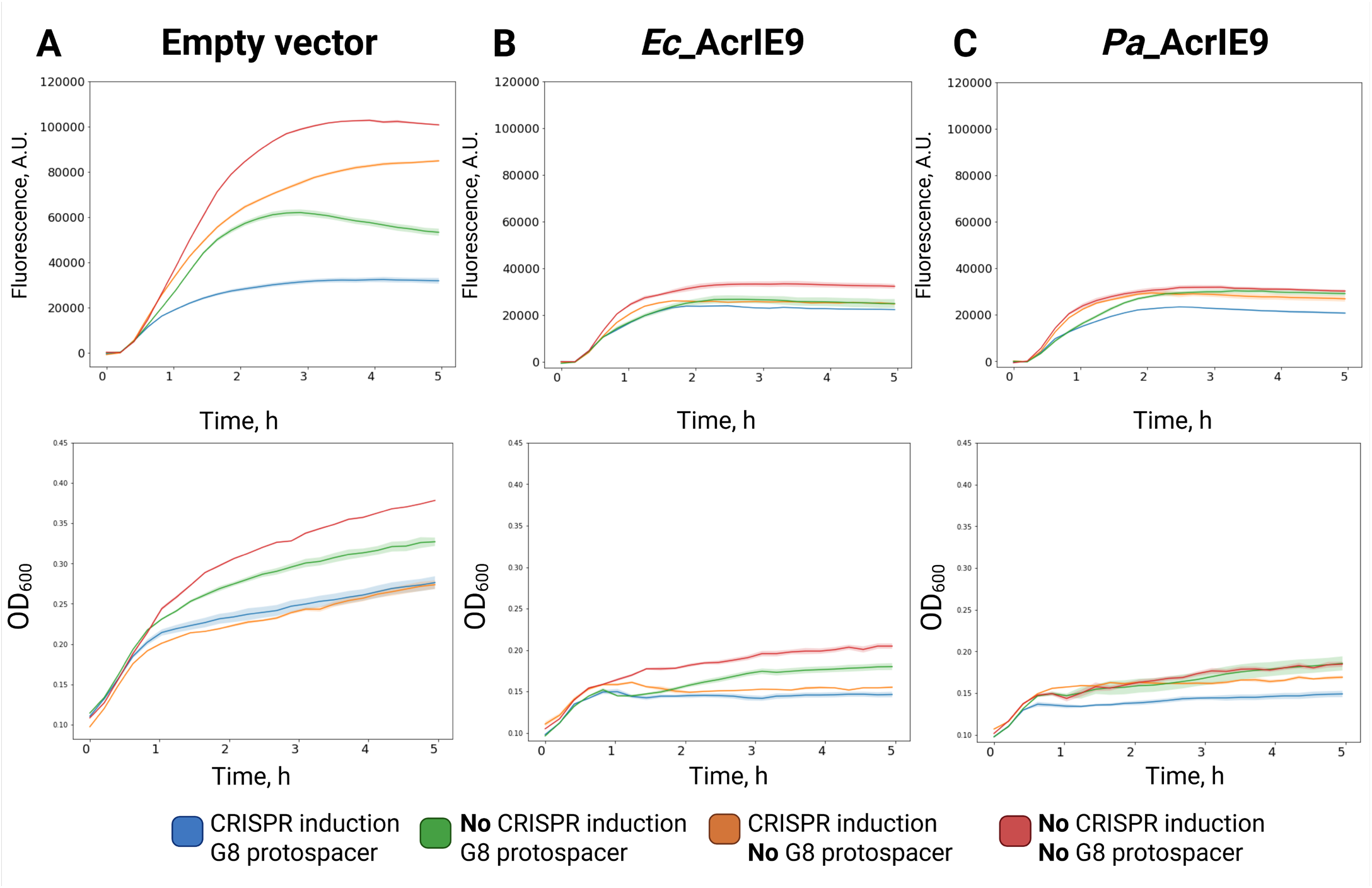
Additional data for Figure 5B. Dynamics of sfGFP production in KD454 cells carrying an empty pBAD vector **(A)** or expressing *Ec*_AcrIE9 **(B)** or *Pa*_AcrIE9 **(C)**. Top - Dynamics of sfGFP production measured via fluorescence accumulation. Bottom – optical density for the respective cultures. Acr expression inhibits *E*.*coli* growth, yet suppresses Cascade-mediated transcriptional silencing.

**Figure S4.**
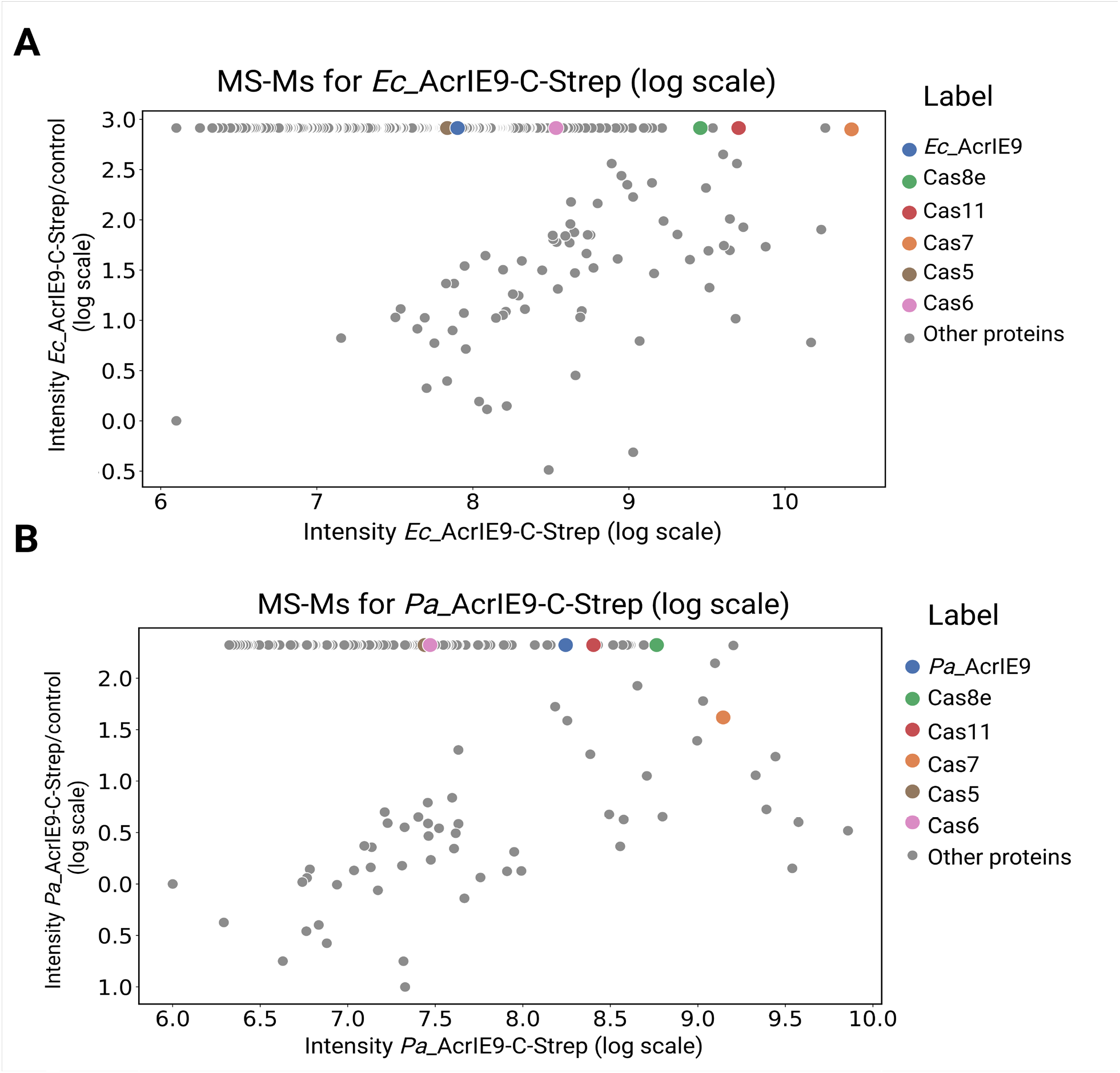
AcrIE9 interacts with Cas7 *in vivo*. Tandem mass-spectrometry analysis of the *in vivo* pull-downs performed with C-Strep *Ec*_AcrIE9 **(A)** or *Pa*_AcrIE9 **(B)** in *E*.*coli* KD263 after induction of CRISPR-Cas expression. Control pull-downs performed with AcrIE9 without Strep tag account for non-specific binding of proteins from the lysate to the column and were used to normalize protein intensity.

**Figure S5.**
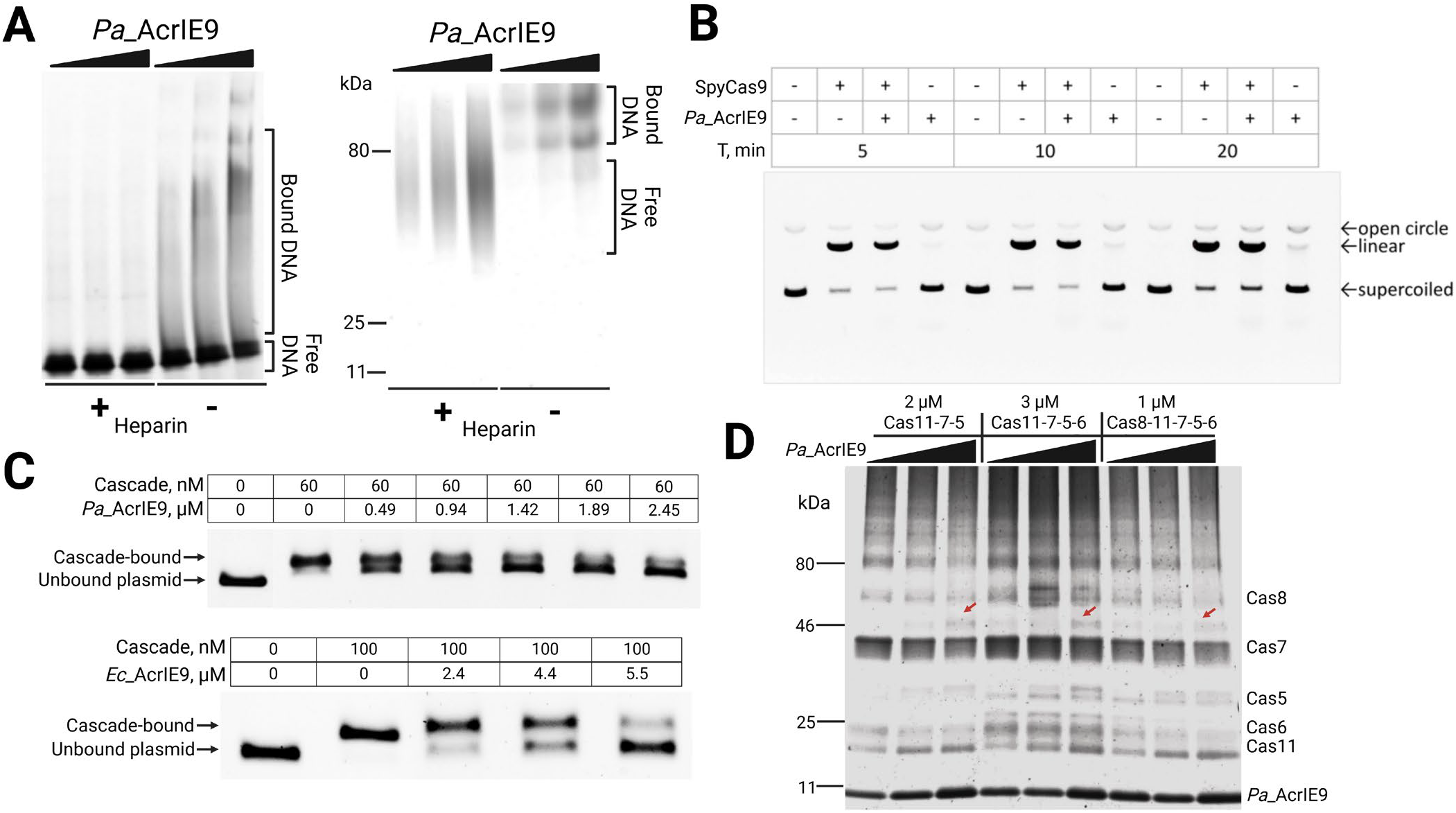
AcrIE9 binds to dsDNA and inhibits Cascade interaction with DNA target. **(A)** EMSA of *Pa*_AcrIE9 with dsDNA performed in conditions of competition with heparin. DNA (left) and protein (right) staining indicates formation of high-molecularweight complexes and reduction of their size in the presence of heparin. Positions of control proteins with known molecular masses are labeled. 0.125 μM of 52-bp dsDNA was incubated with increasing concentrations of *Pa*_AcrIE9 (26, 46 and 92 μM). **(B)** Target plasmid DNA (1.2 nM) cleavage with 75 nM of SpyCas9 in the presence of 14 μM AcrIE9. **(C)** EMSA with a fixed concentration of *Ec*_IE Cascade and increasing concentration of AcrIE9 demonstrates disruption of *Ec*_IE Cascade:DNA complexes. Top panel: 60 nM of *Ec*_IE Cascade has been incubated with 4.5nM of target plasmid and *Pa*_AcrIE9. Bottom panel: 100 nM of *Ec*_IE Cascade has been incubated with 1 nM of target plasmid and *Ec*_AcrIE9. **(D)** *In vitro* glutaraldehyde (0.5%) crosslinking assay between *Pa*_AcrIE9 and subassemblies of Cascade complexes with different subunits composition. Each complex was incubated with increasing concentrations of *Pa*_AcrIE9 (5,14,19 μM), resulting in the appearance of novel band migrating above Cas7 (red arrows), that was confirmed to represent Cas7-AcrIE9 adduct.

**Figure S6.**
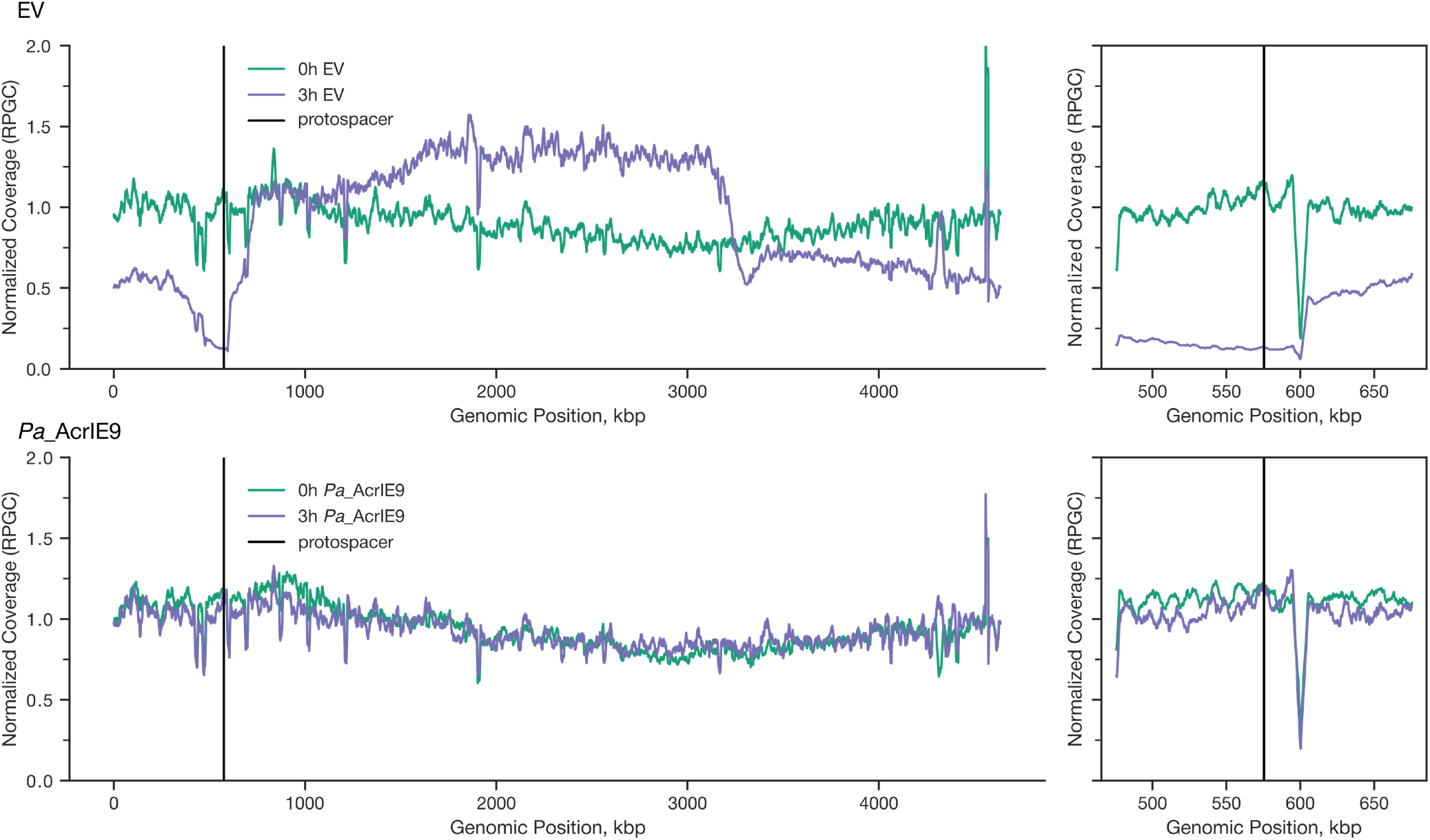
Additional data for Figure 7C. Genomic DNA coverage in self-targeting KD504 cells after CRISPR-Cas induction in conditions of *Pa*_AcrIE9 expression or in the presence of the empty pBAD vector. Bold curves represent moving average for coverage ratios before and 3h after induction of CRISPR-Cas expression. Window size is 10 kbp for the left panel and 5 kbp for the right panel zoomed in at the *yihN* protospacer region (black vertical line).

**Figure S7.**
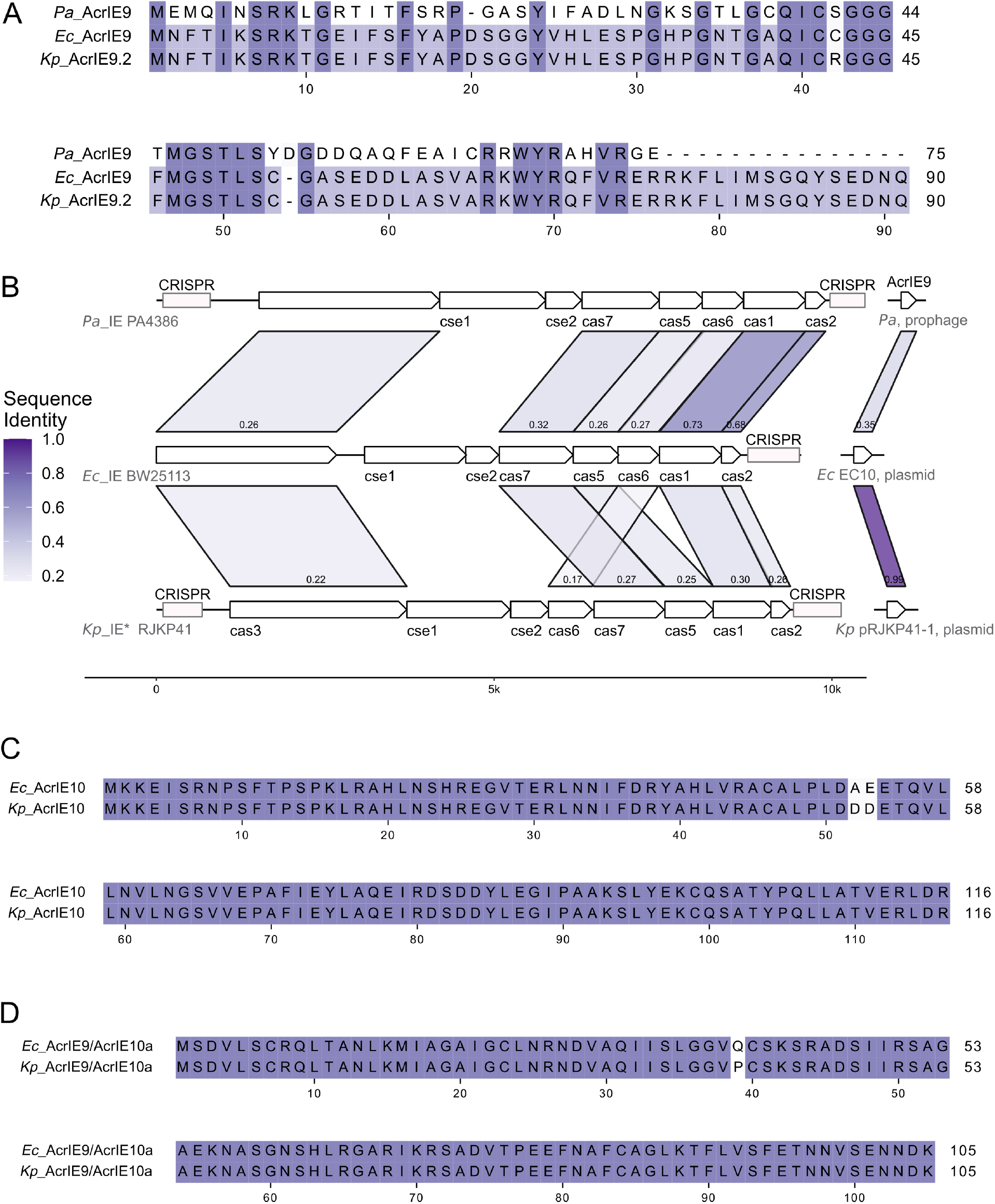
Different Type I-E CRISPR-Cas systems are inhibited by AcrIE9 proteins. **(A)** Multiple sequence alignment of experimentally studied AcrIE9 proteins. **(B)** Sequence identity between CRISPR-Cas systems and corresponding AcrIE9 proteins. Color represents the identity level (shown at the bottom on each colored segment). Sequence identity less than 0.2 is not shown. **(C)** Sequence alignment between AcrIE10 from *E. coli* and *K. pneumoniae*. **(D)** Sequence alignment between AcrIE9/10a from *E. coli* and *K. pneumoniae*. Pa – *Pseudomonas aeruginos*a, Ec – *Escherichia coli*, Kp – *Klebsiella pneumoniae*.

**Figure S8.**
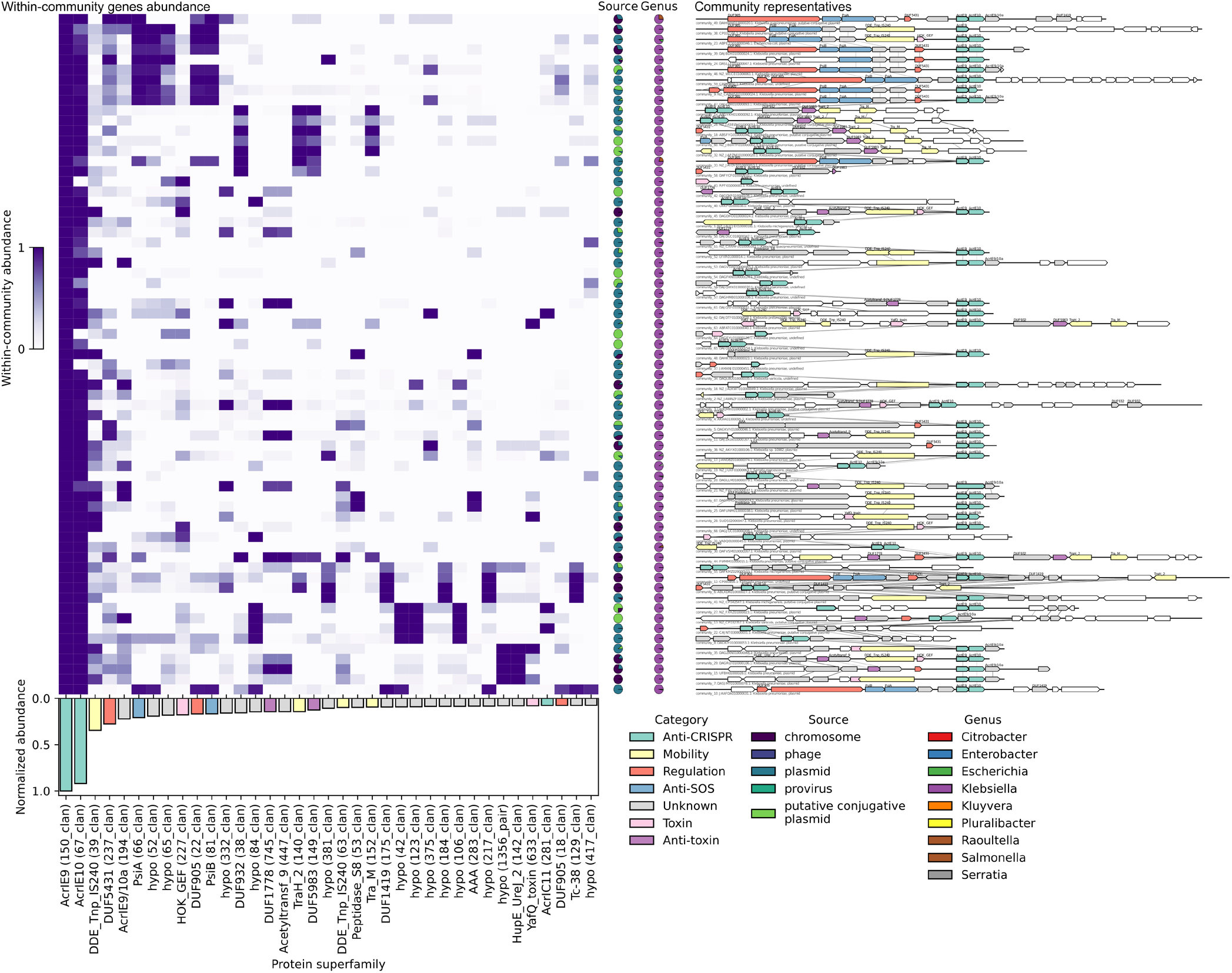
The diversity of loci, encoding AcrIE9. **(Left upper)**, within-community abundance of the most common protein superfamilies, encoded in different communities of loci, encoding AcrIE9. The color of each cell indicates the fraction of loci in the community (rows), encoding the corresponding protein superfamily (columns). Pie-charts indicate the localization (source) and taxonomy (genus) content of a community. **(Left bottom)**, community-normalized abundance of proteins for the most common proteins, encoded among all loci (normalized abundance more than 0.075, See `Loci diversity analysis` in `Methods` for details). The bars are colored according to a functional category that has been assigned manually. **(Right)** Representative genomic loci of the corresponding communities. Shown is the locus with the highest degree of centrality within the given community. Non-compressed variant of the Figure S8 can be found as an **Extended Data File 2**.

**Figure S9.**
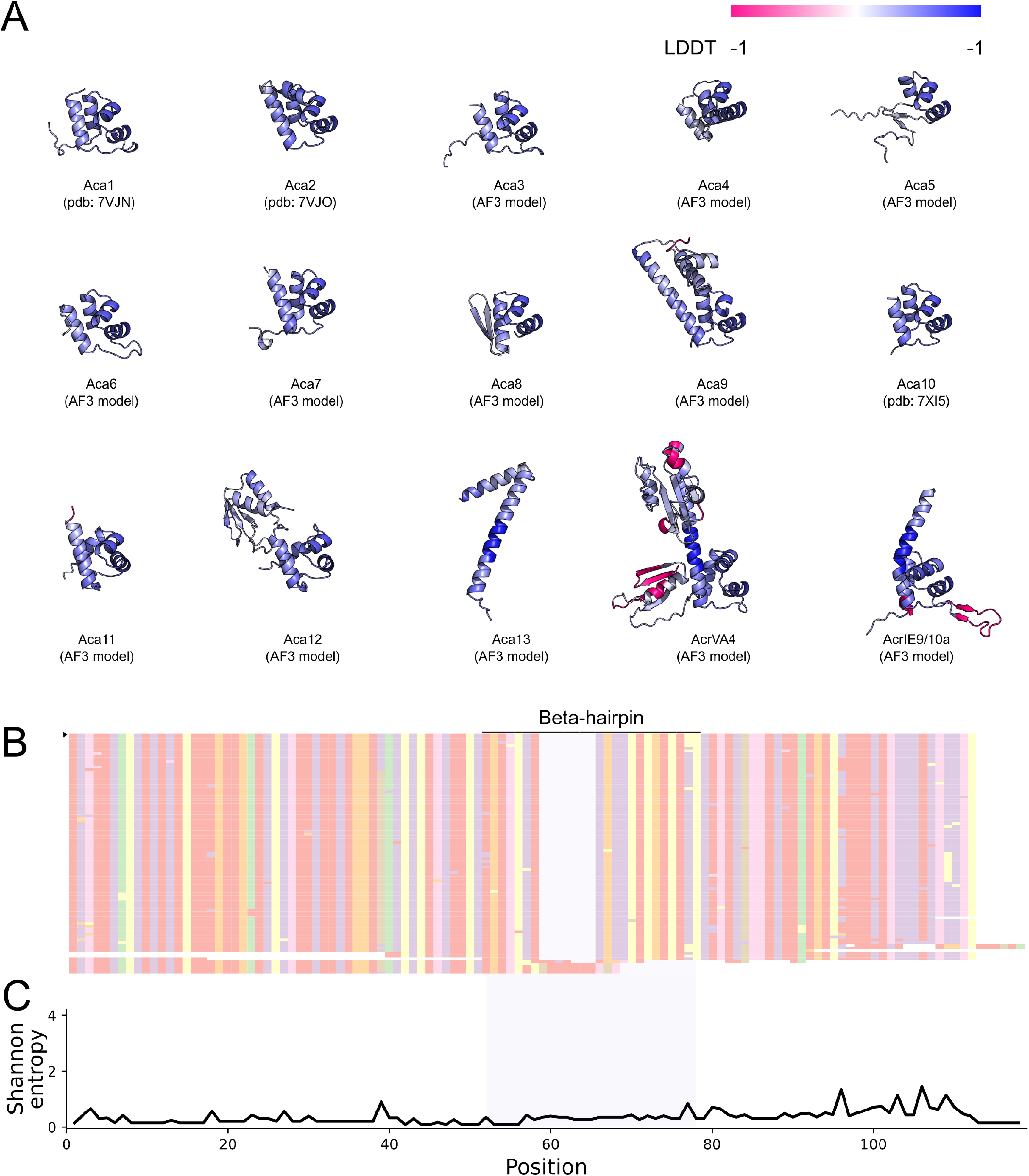
AcrIE9/10a is predicted to be structurally similar to Aca proteins and is characterized by insertion of β-hairpin. **(A)** Comparison of protein structures for previously discovered Aca proteins. Multiple sequence alignment of protein structures was performed with FoldMason WebServer^123^. Protein chains are colored according to the obtained structure-based score (LDDT) of a MSA, where -1 represents unaligned regions. All used protein sequences are listed in **Supplementary Table 3. (B)** Multiple sequence alignment of unique protein sequences belonging to AcrIE9/10a protein superfamily. AcrIE9/10a from AcrIE9 locus studied in this paper is marked with a triangle. The amino acids color scheme is adapted from Clustal color scheme. Residues are hidden for simplicity. **(C)** Shannon entropy calculated to 21-letters alphabet (gaps counted) for MSA on **(B)**.

**Figure S10.**
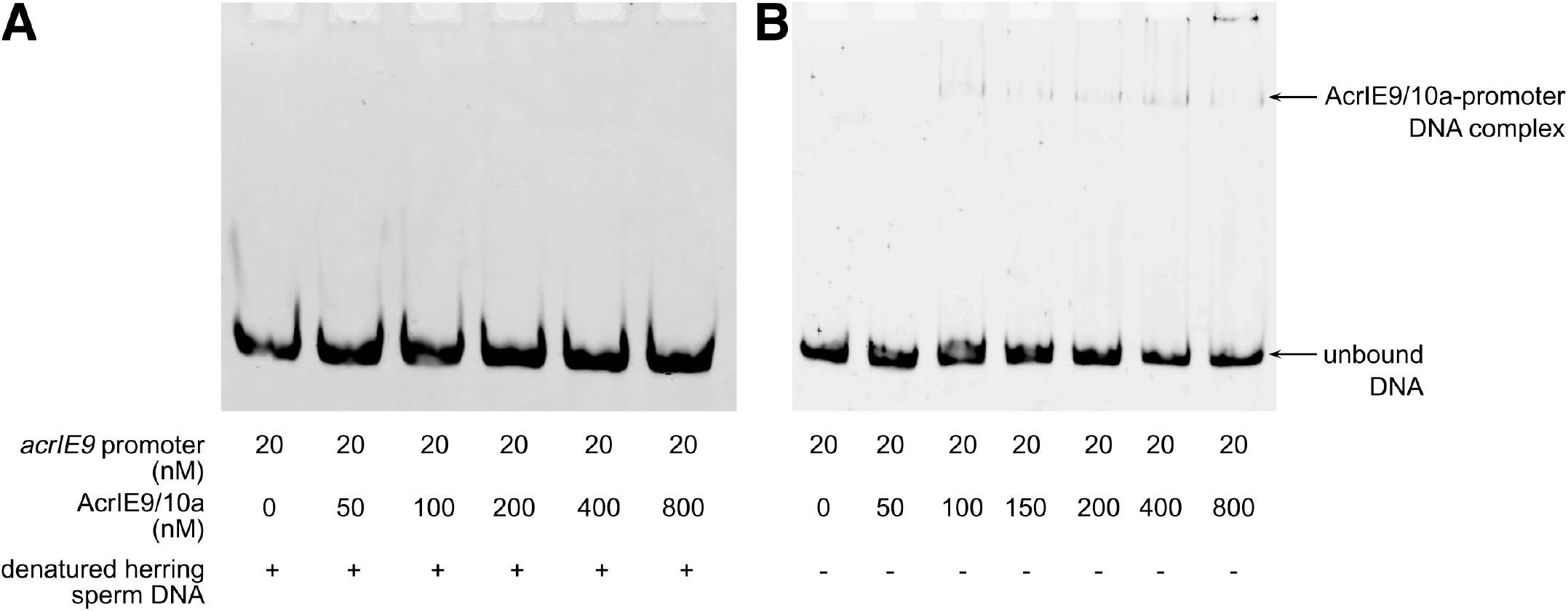
AcrIE9/10a has non-specific DNA binding activity. EMSA of *Kp*_AcrIE9/10a with 300 bp 5-FAM labeled dsDNA containing *acrIE9* promoter performed in conditions of competition with an excess of non-specific DNA **(A)** or without competition **(B)**.

**Supplementary Table 1.**
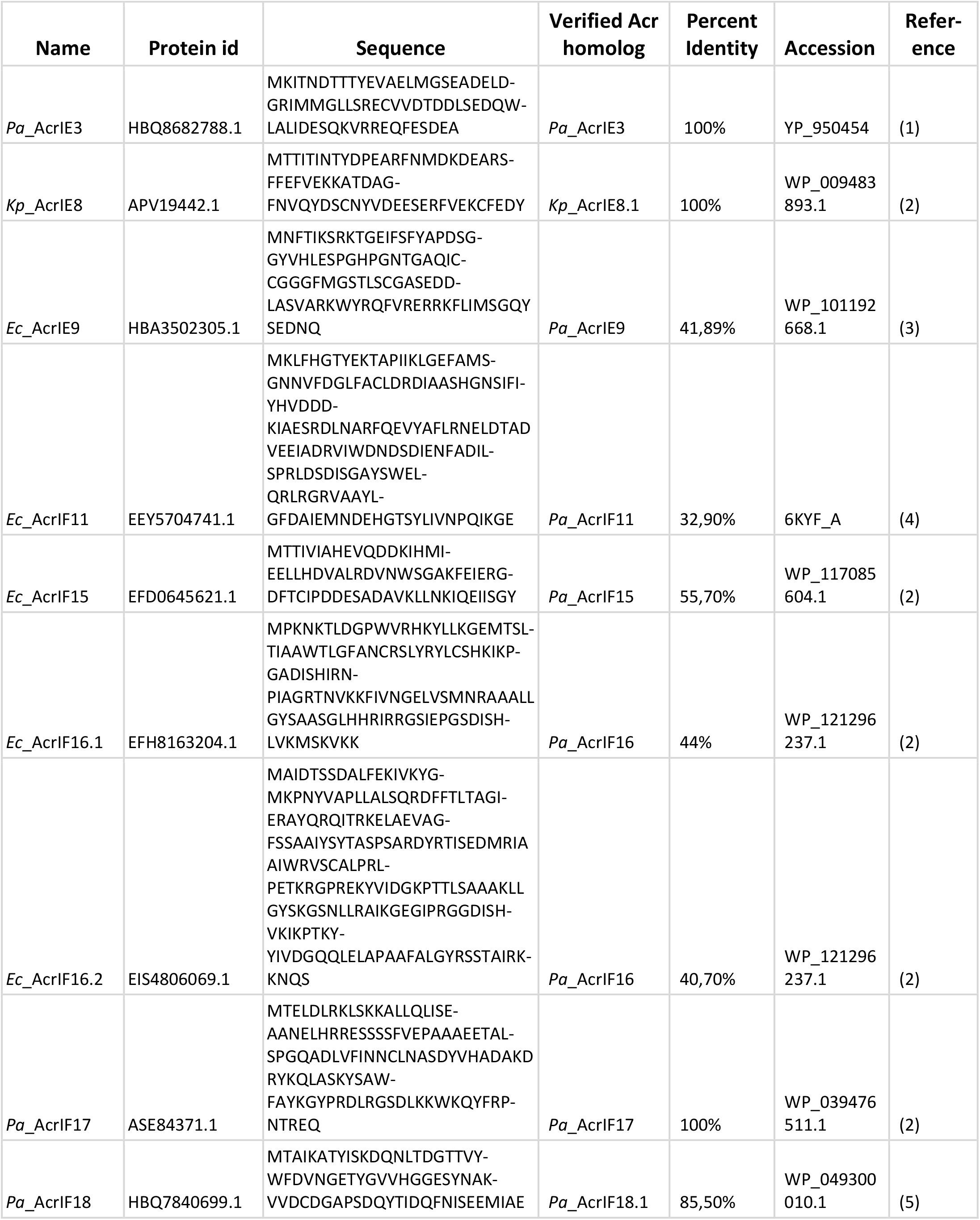
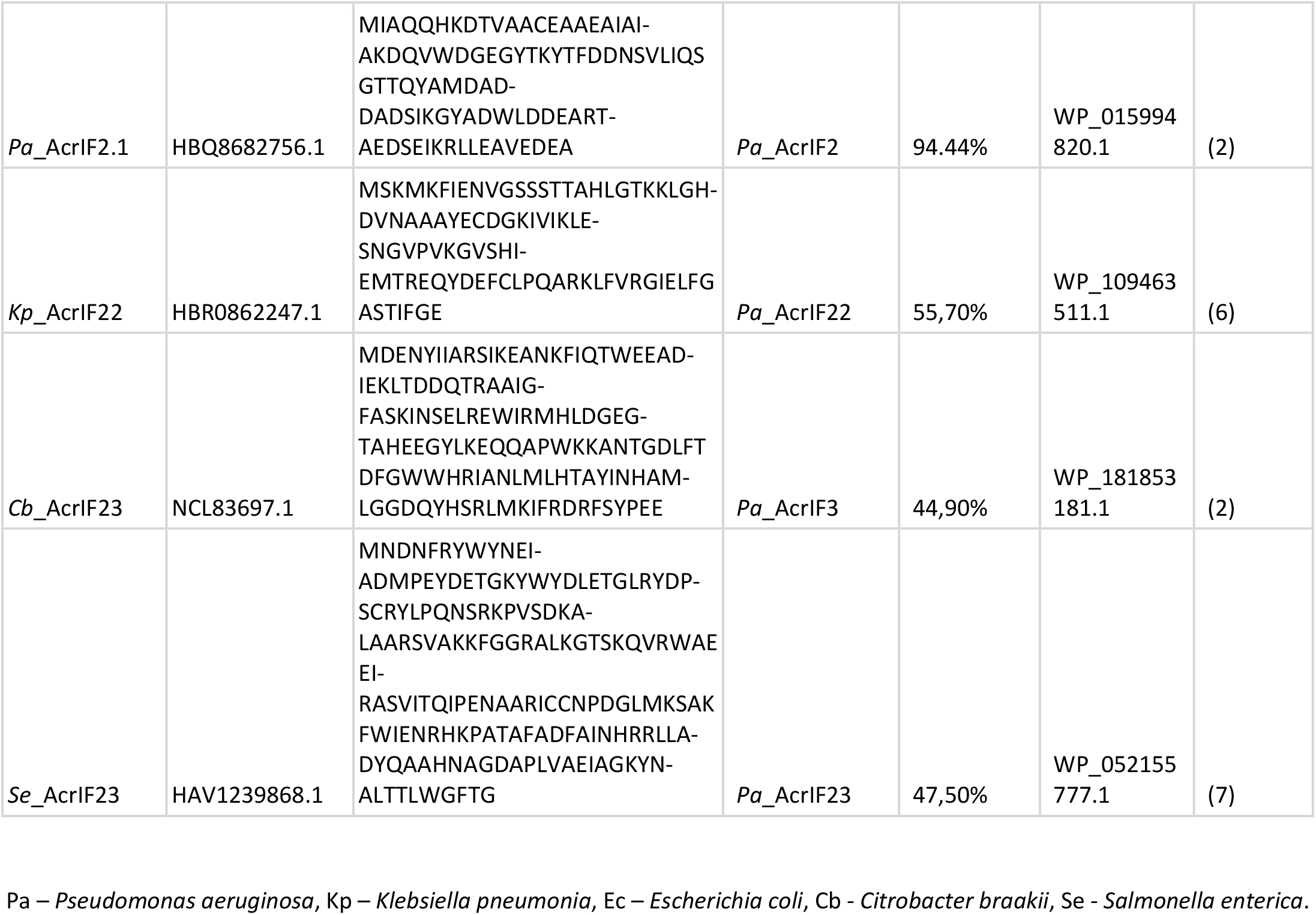
Dataset of Acr homologs studied in this work.

**Supplementary Table 2.**
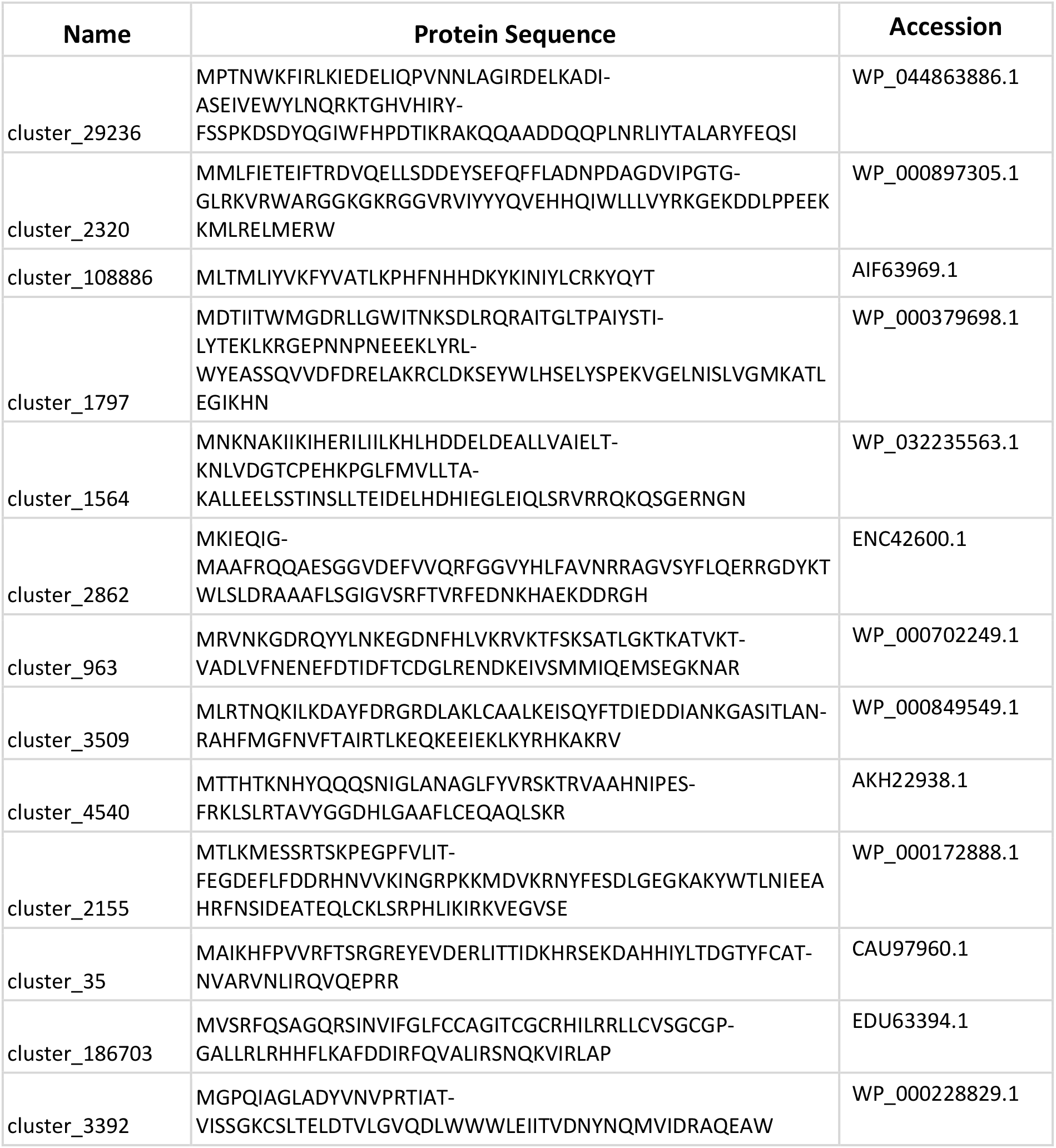
Dataset of ML-predicted Acr candidates studied in this work.

**Supplementary Table 3.** Aca proteins analyzed in this work.

| Protein | Accession | Source | Reference |
| --- | --- | --- | --- |
| Aca1 | 7VJN | PDB | (8) |
| Aca2 | 7VJO | PDB | (8) |
| Aca3 | WP_049360086.1 | NCBI Protein | (9) |
| Aca4 | WP_079381596.1 | NCBI Protein | (9) |
| Aca5 | WP_074032234.1 | NCBI Protein | (9) |
| Aca6 | WP_035450933.1 | NCBI Protein | (9) |
| Aca7 | WP_064700810.1 | NCBI Protein | (9) |
| Aca8 | YP_009272953.1 | NCBI Protein | (9) |
| Aca9 | WP_131138857.1 | NCBI Protein | (9) |
| Aca10 | 7XI5 | PDB | (10) |
| Aca11 | WP_009730540.1 | NCBI Protein | (11) |
| Aca12 | WP_033585136.1 | NCBI Protein | (11) |
| Aca13 | WP_094140901.1 | NCBI Protein | (11) |
| AcrVA4 | WP_046699156.1 | NCBI Protein | (12) |
| AcrIE9/10a | WP_013023810.1 | NCBI Protein |  |

**Supplementary Table 4.**
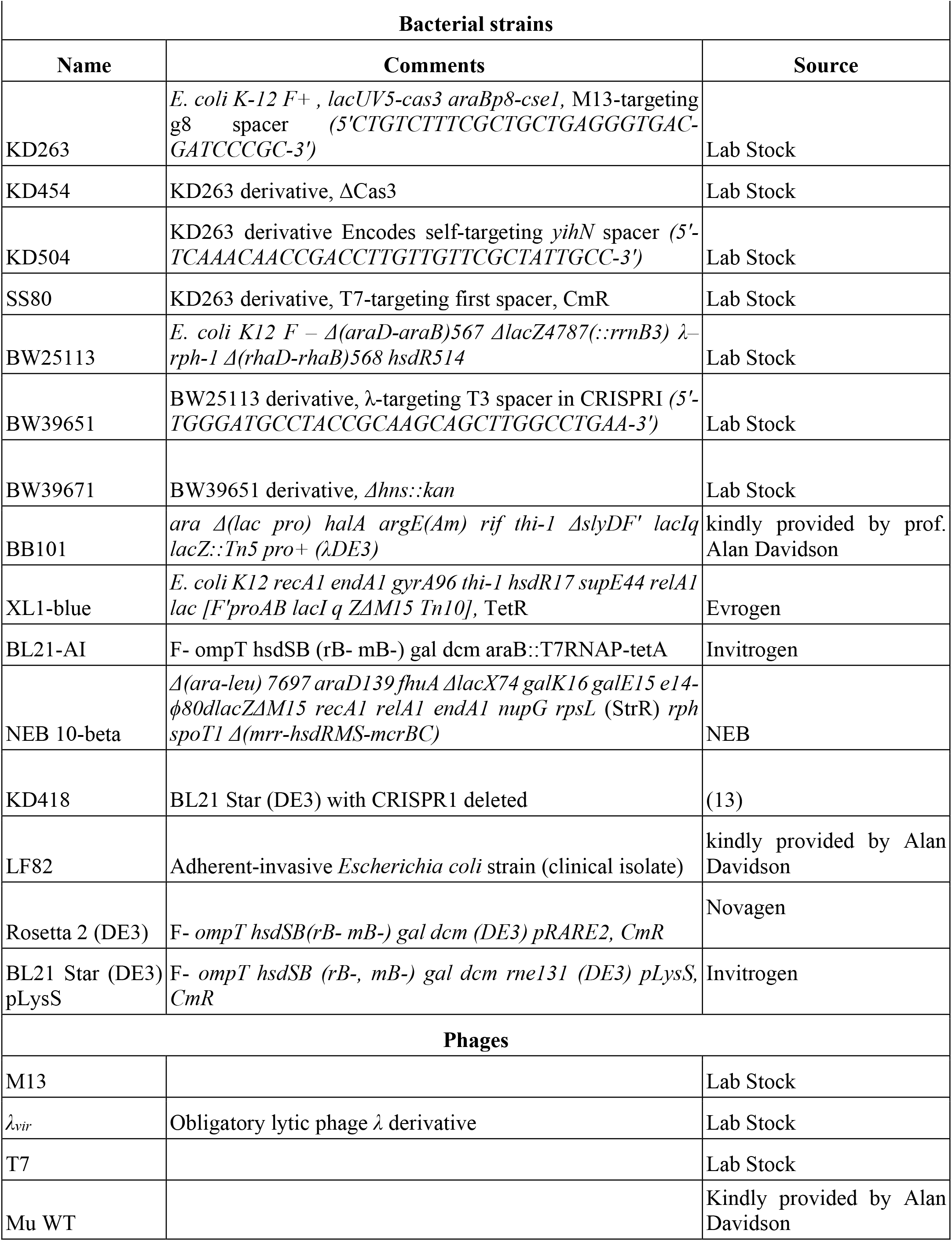
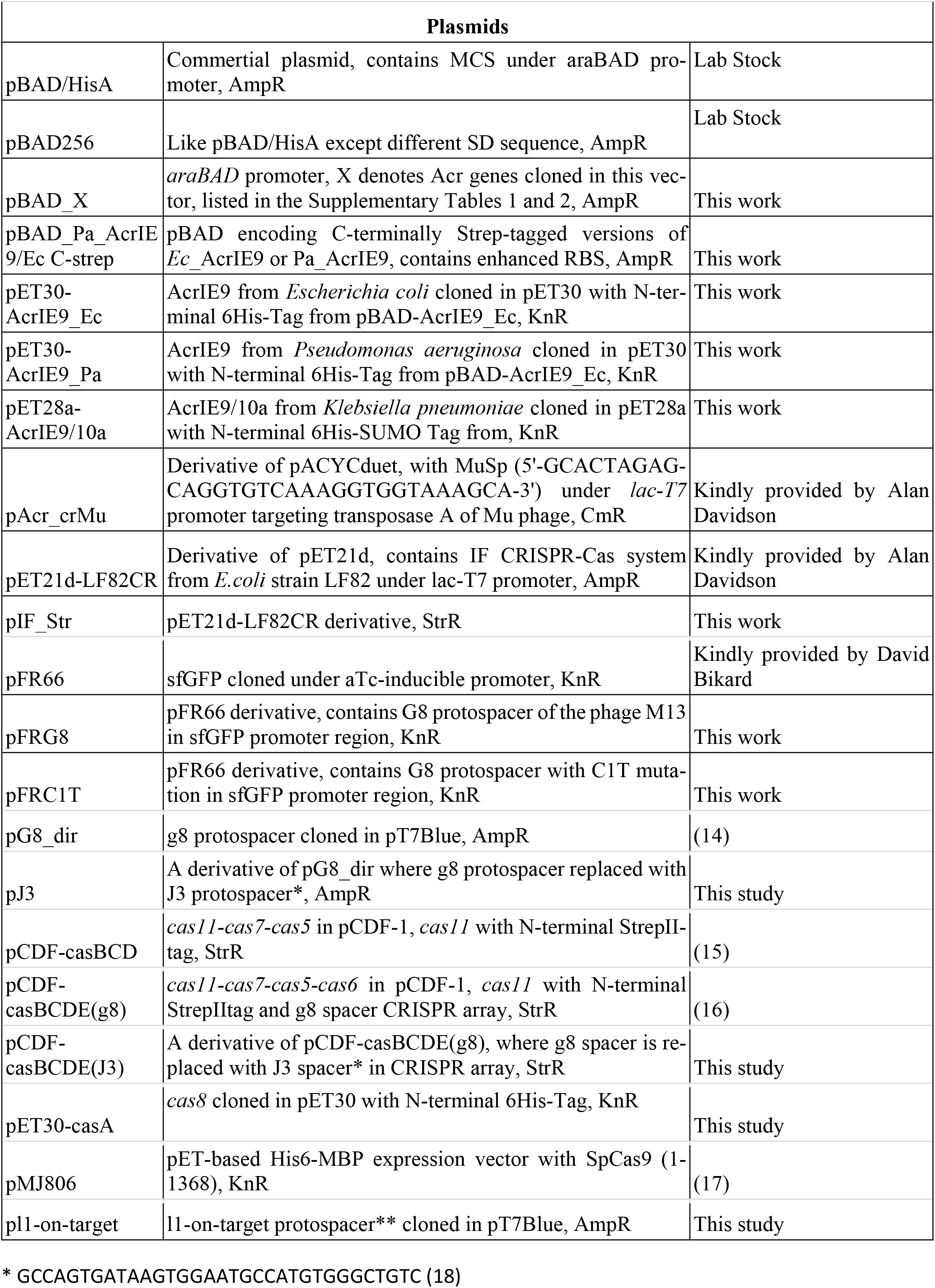
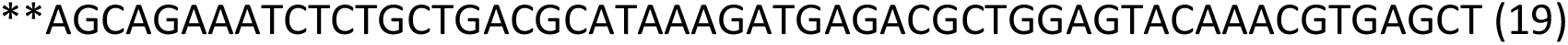
List of strains, phages and plasmids used in this work.

**Supplementary Table 5.**
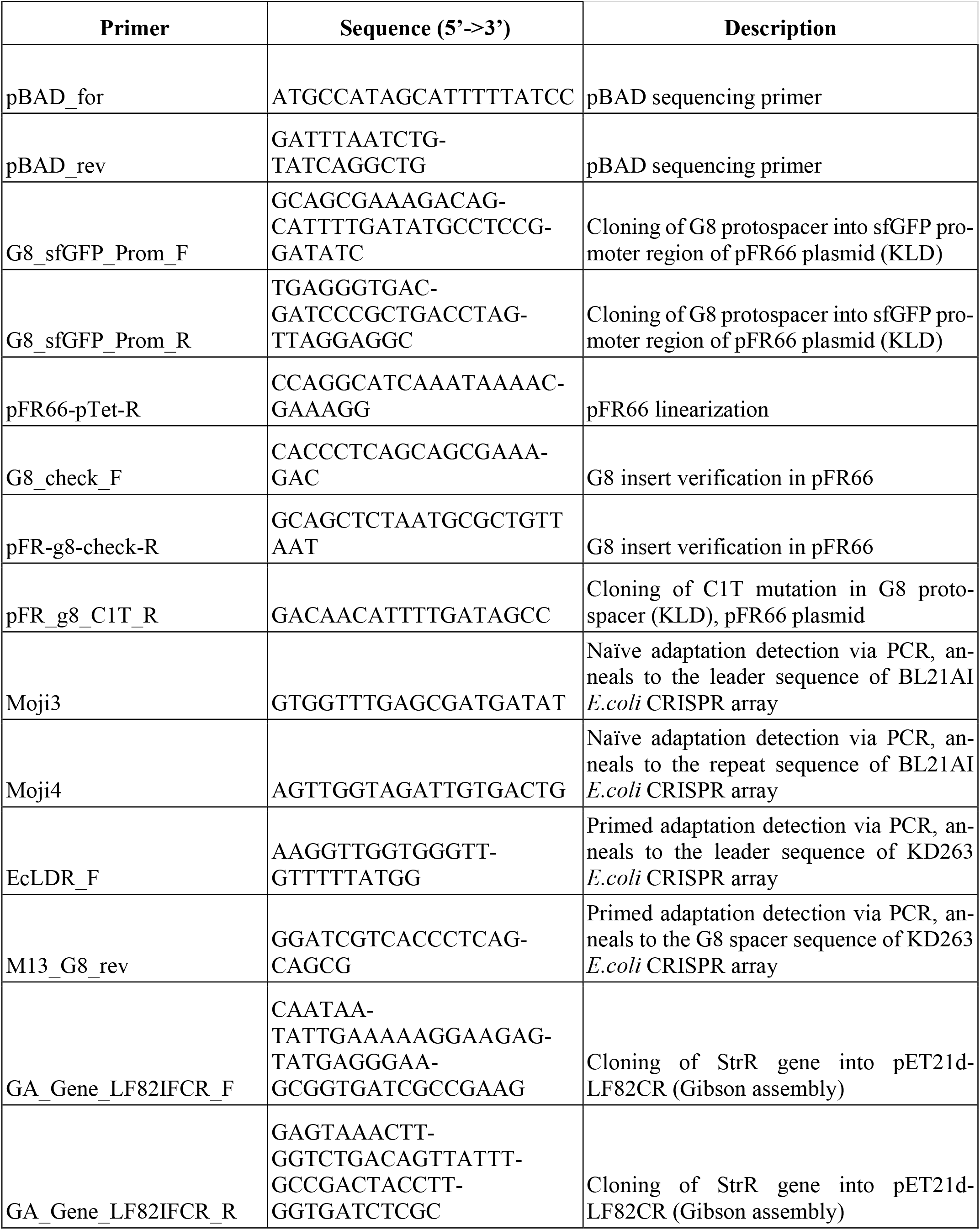
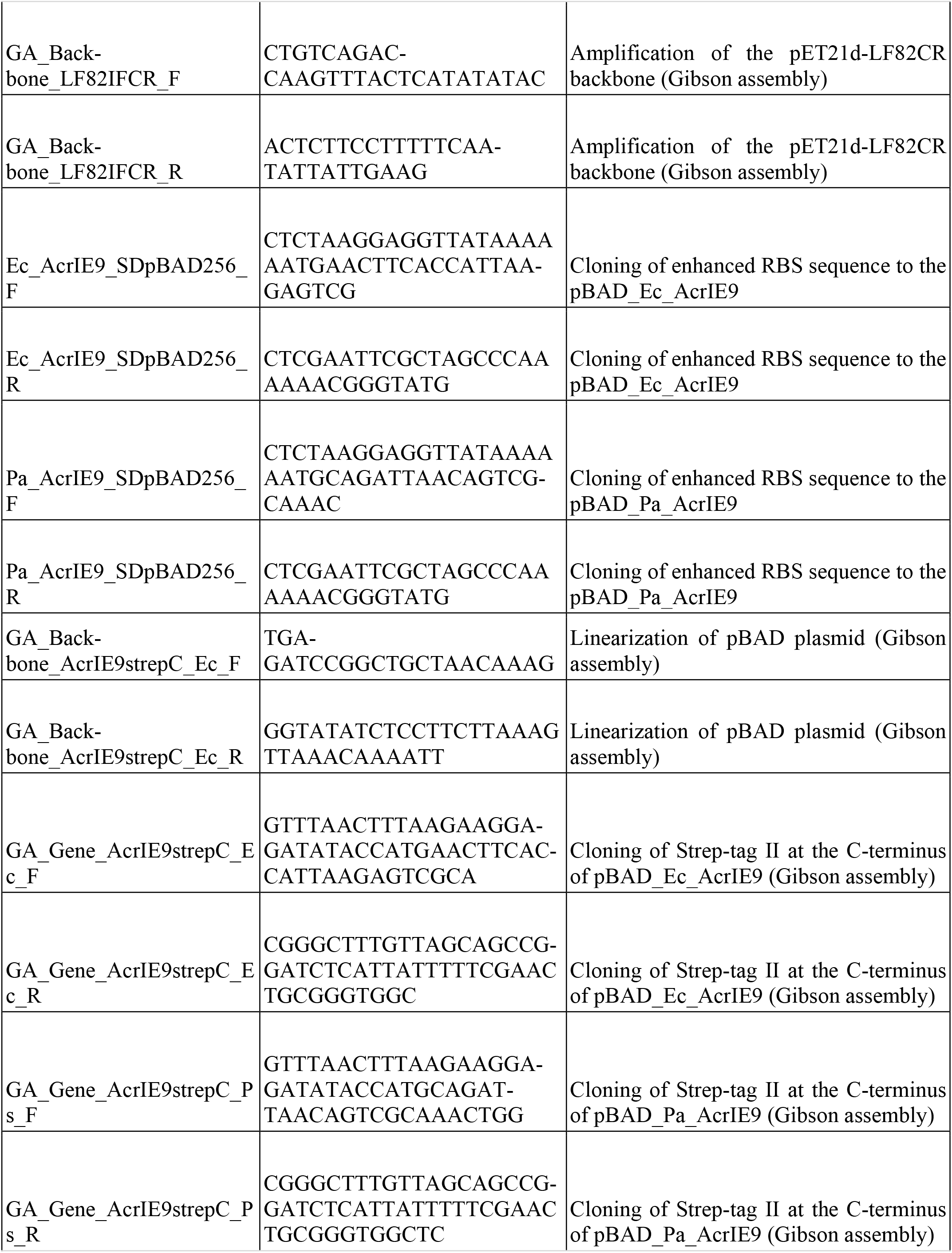
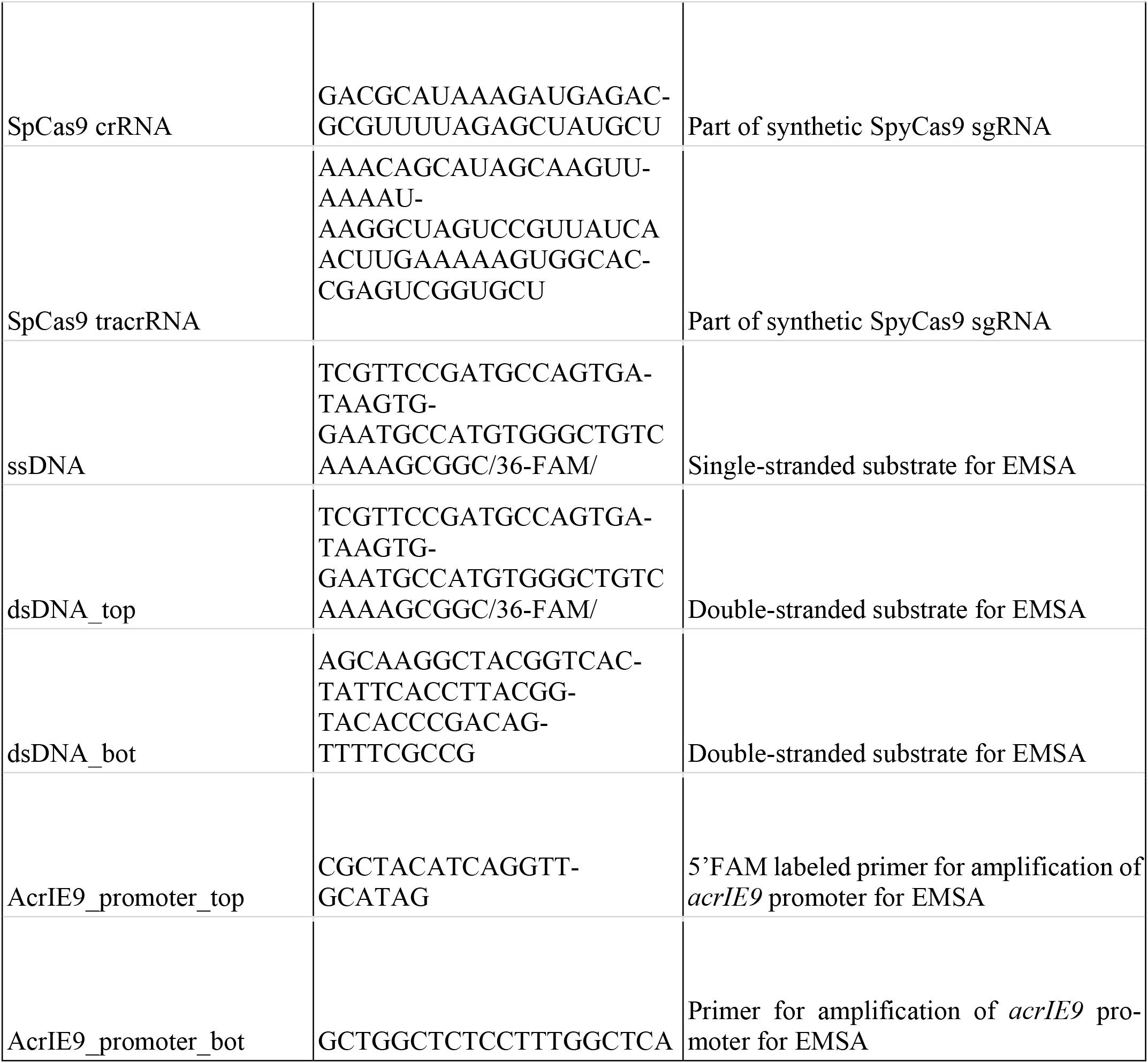
List of primers used in this work.

