## Supplementary figures and images for "A census of anti-CRISPR proteins reveals AcrIE9 as an inhibitor of *Escherichia coli* K12 Type IE CRISPR-Cas system"

### Extended Data File 2 - AcrIE9 encoding loci.pdf

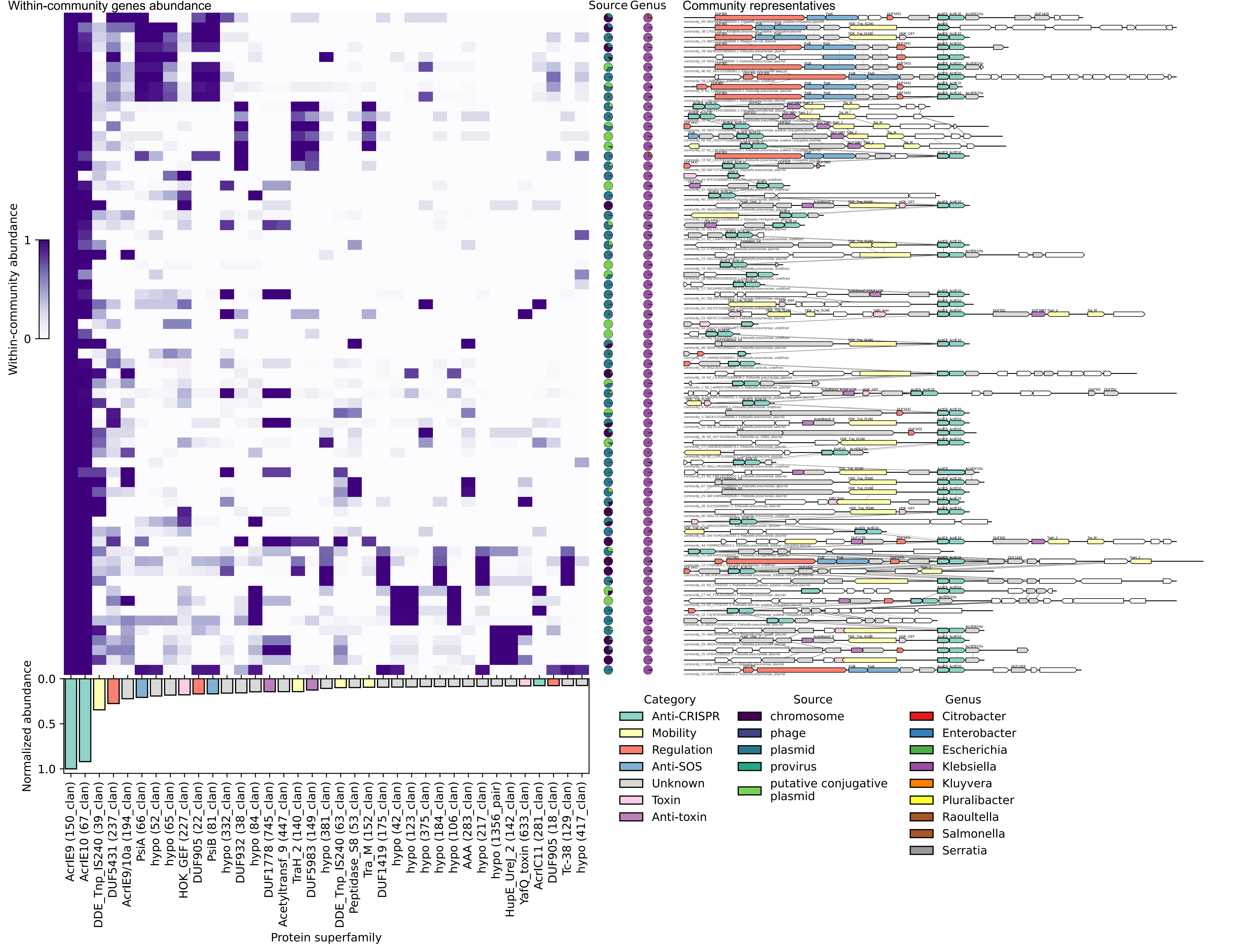
